# An OmpX mRNA-LNP vaccine induces protective humoral and cellular immunity against *Klebsiella pneumoniae* in a C57BL/6 mouse model

**DOI:** 10.64898/2026.10.26.761762

**Authors:** Yunwang Zhao, Hongxian Cheng, Shiyi Zhou, Lei He, Chenlin Xiu, Baihai Huang, Xiongxiong Lu

## Abstract

*Klebsiella pneumoniae* (KP) poses a significant threat to global public health, and effective preventive measures are urgently needed. The widespread antibiotic resistance of KP underscores the critical need for vaccine-based strategies. Although numerous vaccine candidates against KP have been investigated, none has yet been licensed. mRNA-LNP vaccines against KP have only recently begun to be explored, and their ability to deliver conserved outer membrane protein antigens remains largely untested. In this study, we constructed an mRNA vaccine encoding the highly conserved outer membrane protein X (OmpX) of KP-the first mRNA-LNP vaccine targeting this antigen-and evaluated its immunogenicity and protective efficacy in mice. Immunoinformatics analyses predicted concordance between the in silico and in vivo outcomes: bioinformatics tools were used to evaluate the antigenicity, immunogenicity, physicochemical parameters, and secondary and tertiary structures of OmpX, and the C-ImmSim server was used to simulate the vaccine-elicited immune responses. We then performed mouse immunization and challenge experiments to characterize the humoral and cellular immune responses induced by the OmpX vaccine. The mRNA vaccine was immunogenic in mice, eliciting high titers of OmpX-specific antibodies. OmpX-Fc mRNA-LNPs also induced robust cell-mediated immunity, with significantly elevated IFN-γ and IL-4 levels indicative of combined Th1/Th2 cellular responses. Serum from OmpX-immunized mice promoted significantly greater complement-mediated KP cell lysis than serum from the alum or luciferase mRNA-LNP control groups. Notably, OmpX-Fc mRNA-LNP immunization conferred complete protection against lethal KP challenge, with 100% survival and a significant reduction in visceral bacterial burden. These data establish a promising mRNA vaccine strategy for inducing broad immune responses and promoting serotype-independent immunity against KP.

**Graphical Abstract:** 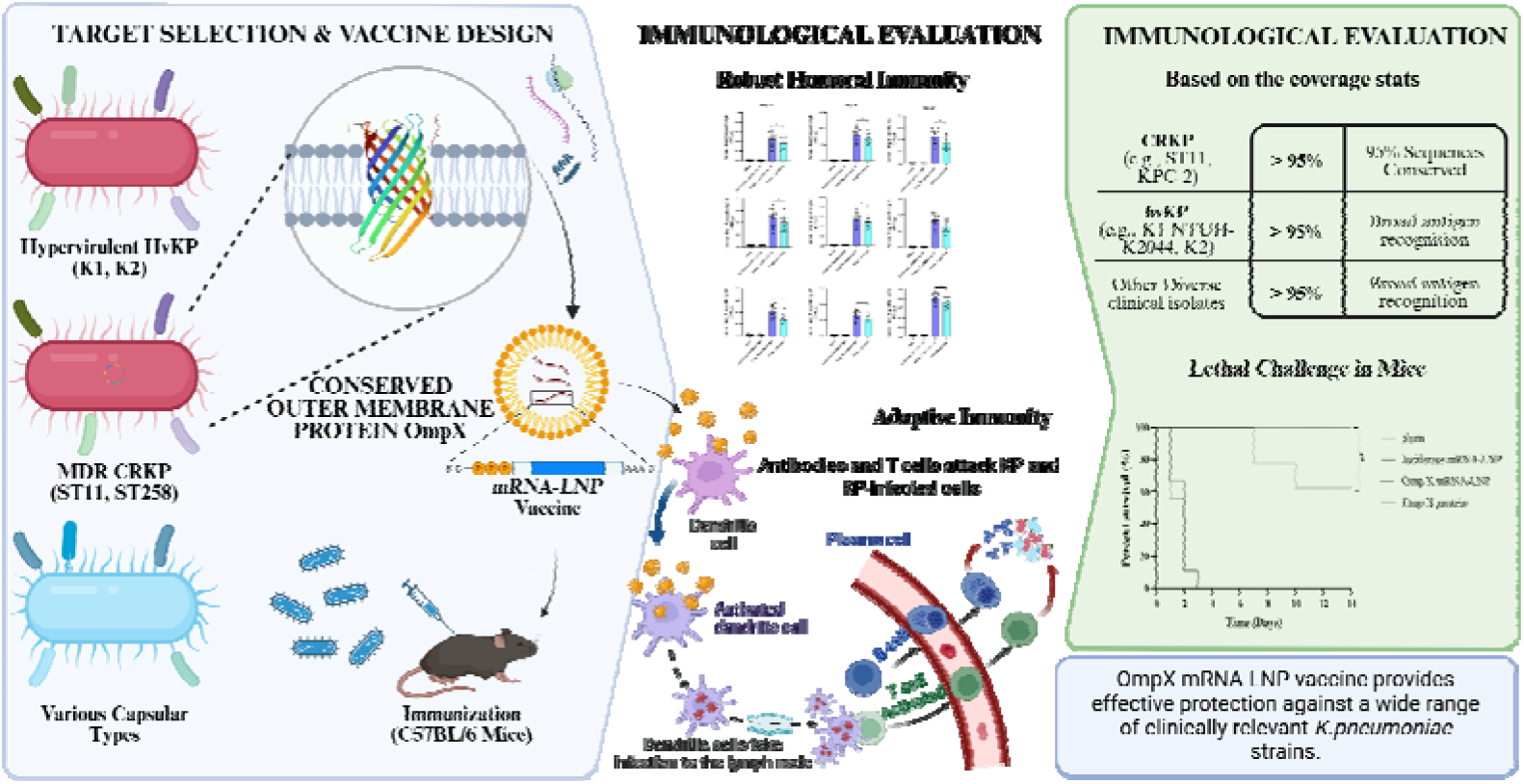

---

*Klebsiella pneumoniae* (KP), a Gram-negative opportunistic pathogen, poses a serious global public health threat because of its escalating antibiotic resistance and clinical severity [1, 2, 28–30]. This pathogen is associated with a broad spectrum of infections, including pneumonia, urinary tract infections, and life-threatening bacteremia and sepsis [3]. Based on genomic and phenotypic profiles, KP strains can be categorized into three clinically significant subtypes: (1) opportunistic strains that cause infections in immunocompromised hosts; (2) hypervirulent strains characterized by extensive virulence-associated gene clusters that facilitate tissue invasion; and (3) multidrug-resistant (MDR) strains harboring plasmid-borne resistance determinants [4]. Although antibiotics remain the primary therapeutic strategy for severe KP infections, their efficacy is increasingly compromised by adverse effects and inappropriate antimicrobial stewardship, leading to the emergence of MDR strains that pose critical challenges in clinical management.

Vaccines represent a critical prophylactic strategy for mitigating KP colonization and infection. Current KP vaccine candidates include whole-cell vaccines and cell lysates, capsular polysaccharide-based vaccines, lipopolysaccharide-targeted approaches, protein subunit vaccines, outer membrane vesicles (OMVs) [5, 33], and glycoconjugate platforms. However, none has achieved clinical licensure, owing to limitations such as high production costs, poor stability, toxicity concerns, and inadequate protective efficacy in preclinical models and human trials. A key obstacle is the antigenic diversity among KP strains, which severely restricts the cross-protective capacity of conventional vaccines. These challenges underscore the urgent need for innovative vaccine platforms capable of eliciting broad-spectrum protection against evolving KP strains.

Compared with subunit or inactivated vaccines against KP, the mRNA-LNP platform offers several practical advantages for this application [38–40]. OmpX is a hydrophobic transmembrane porin that is difficult to express and purify in recombinant form from *E. coli*-a major bottleneck for protein subunit approaches. Encapsulating OmpX-encoding mRNA within LNPs bypasses this challenge entirely by enabling in situ antigen production within host cells after translation of the delivered transcript. Moreover, the modular architecture of mRNA allowed us to append a human IgG Fc tag to the OmpX coding sequence, enhancing antigen presentation to follicular dendritic cells and improving germinal center responses. The LNP formulation (lipid01/DSPC/cholesterol/DMG-PEG2k) also serves as an intrinsic adjuvant by activating the STING and TLR4 pathways [41, 42], promoting the Th1/Th17 polarization required for mucosal anti-KP immunity. This approach has demonstrated success across diverse infectious disease models, including bacterial (Mycobacterium tuberculosis) and parasitic (Plasmodium) pathogens [46, 47]. Notably, Iwanaga et al. demonstrated that immunization with recombinant OmpX combined with a mucosal Th17 adjuvant elicits lung-resident memory T cells that protect against heterologous KP challenge [31, 32]; however, whether OmpX can be effectively delivered by an mRNA-LNP platform to elicit systemic humoral and cellular immunity has not been investigated.

In parallel, the rapid advancement of computational tools and immunoinformatics has accelerated reverse vaccinology [6]. Immunoinformatics is now widely employed for epitope discovery, precision immune engineering, and accelerated vaccine design, enabling the screening of candidate antigens with protective potential at the whole-genome level. This strategy is convenient, widely applicable, safe, labor-saving, and cost-effective. However, the concordance between immunoinformatics predictions and in vivo experimental outcomes for OmpX-based vaccines has not been systematically evaluated.

In this study, we designed an mRNA vaccine encoding OmpX and evaluated its potential against KP infection. Bioinformatics tools were used to predict the antigenicity, immunogenicity, secondary structure, and physicochemical properties of OmpX. We assessed the protective potential of the vaccine through tertiary structure modeling, molecular docking, and in silico immunostimulation predictions. Finally, animal experiments were conducted to validate the immunological properties of the OmpX vaccine. The vaccine maximized adaptive immune responses in a mouse model, inducing specific humoral and cell-mediated immunity. The utility of bioinformatics in reverse vaccinology was also assessed by comparing computational predictions with experimental data.

Although a recent study reported an mRNA-LNP vaccine encoding the YidR protein with a tPA signal sequence that conferred Th1-biased protection against KP lung infection [48], the present study extends this platform to OmpX, a conserved β-barrel porin, and demonstrates Fc-tag-enhanced immunogenicity, complement-dependent bactericidal activity, and 100% survival against lethal intraperitoneal challenge.

## Materials and Methods

### Bioinformatics prediction

#### OmpX protein sequence retrieval

The amino acid sequence of OmpX (UniProt accession number A0A486PWV9) was retrieved from UniProt (http://www.uniprot.org).

#### Immunogenicity, antigenicity, allergenicity, and toxicity prediction

Immunogenicity and antigenicity are critical determinants of vaccine efficacy against KP infection. To assess immunogenicity, we submitted the OmpX amino acid sequence to the IEDB immunogenicity tool (http://tools.iedb.org/immunogenicity/) [7, 44], which outputs a score reflecting the probability of T-cell epitope induction. For antigenicity evaluation, we used the Kolaskar-Tongaonkar antigenicity method (EMBOSS antigenic; http://imed.med.ucm.es/Tools/antigenic.pl), which estimates protective antigen potential through sequence-based machine learning.

The AllerTOP v.2.0 (http://www.ddg-pharmfac.net/AllerTOP/) and AllergenFP v.1.0 (http://ddg-pharmfac.net/AllergenFP/) servers were used to assess allergenicity [8, 45], and the ToxinPred server (http://crdd.osdd.net/raghava/toxinpred/) was used to predict toxicity. AllerTOP v.2.0 and ToxinPred results were reported as sensitizing or non-sensitizing and toxic or non-toxic, respectively.

#### Physicochemical properties and solubility

Sequence-derived physicochemical features of OmpX-theoretical pI, predicted intracellular/extracellular half-life, instability index, and aliphatic index-were computed using the ExPASy ProtParam pipeline (https://web.expasy.org/protparam/) [9], which derives these parameters directly from the primary amino acid composition. Vaccine solubility was predicted using the Protein-Sol server (https://protein-sol.manchester.ac.uk/), a sequence-based method in which scores range from 0 to 1 and values exceeding 0.45 indicate good solubility.

#### Secondary and three-dimensional structure prediction

OmpX secondary structure elements (α-helices, β-strands, and random coils) were predicted by PSIPRED [10] using position-specific scoring matrices. For tertiary structure modeling, the 3Dpro server was queried to generate an initial all-atom model via a fragment-assembly approach, which was subsequently refined using GalaxyRefine [11] to optimize backbone geometry and rotamer conformations.

#### Tertiary structure validation and molecular docking

The ERRAT server (https://saves.mbi.ucla.edu/) was used to assess the structural quality of the OmpX 3D model.

Refined OmpX PDB files were obtained from the GalaxyRefine web server. PDB files for human leukocyte antigen HLA-DRB1*01:01 (ID: 1AQD), toll-like receptor 2 (TLR-2, 6NIG), and toll-like receptor 4 (TLR-4, 4G8A) were obtained from the Molecular Modeling Database (MMDB, https://www.ncbi.nlm.nih.gov/Structure/).

Ligand-receptor docking analysis was performed using ZDOCK 3.0.2 to evaluate the binding affinity between OmpX and HLA-DRB1*01:01, TLR-2, and TLR-4. Visualization was performed using PyMOL 2.5.3 [12, 13].

#### Prediction of conformational B-cell epitopes

Conformational B-cell epitopes of the OmpX vaccine were predicted using the ElliPro server (http://tools.iedb.org/ellipro/) [14]. These epitopes arise from protein folding, which brings distant residues into spatial proximity, and account for more than 90% of B-cell epitopes.

#### Immune simulation

Immune responses following OmpX vaccine administration were simulated using the C-ImmSim server [15, 44] (https://150.146.2.1/C-IMMSIM/index.php) to predict T-cell, B-cell, and antibody responses. Simulation parameters followed server recommendations: random seed, default volume, and HLA alleles. The simulation was run for 168 time steps, with three OmpX vaccine injections administered at days 0, 14, and 28.

#### Genomic data collection, conservation, and coverage analysis

To evaluate the conservation and coverage of OmpX across KP strains, OmpX amino acid sequences were retrieved from the NCBI GenBank and UniProt databases, encompassing classical (cKP), hypervirulent (hvKP), and multidrug-resistant (MDR) strains. Multiple sequence alignment was performed using Clustal Omega to assess amino acid conservation across clinical subtypes. Epitope region conservation was analyzed by mapping the predicted B-cell and T-cell epitopes onto the multiple sequence alignment, and the percent identity of epitope-containing regions relative to the reference OmpX sequence was calculated.

#### Geographic distribution and subtype coverage mapping

KP clinical strain metadata, including geographic origin (continent and country), isolation source, and clinical subtype classification, were retrieved from public databases. The proportion of strains harboring conserved OmpX epitope sequences was calculated for each geographic region to estimate vaccine coverage. Geographic distribution maps were generated using the ggplot2 package in R (v4.x). Regional subtype distribution was further stratified by continent to visualize the relative prevalence of cKP, hvKP, and MDR strains across Asia, Europe, North America, Africa, and Oceania.

### Experimental validation

#### Bacterial strain and growth conditions

KP strain ATCC 43816 was obtained from the Henan Provincial Industrial Microbial Strain Engineering Technology Research Center (BNCC, China) and cultured in LB broth at 37°C. HEK 293T cells were preserved in 10% FBS-DMEM medium containing 10% DMSO in liquid nitrogen and subsequently cultured in 10% FBS-DMEM medium at 37°C in a 5% CO incubator.

#### Plasmid linearization and *in vitro* transcription

The recombinant plasmid pIVTRup SP-OmpX-Fc-Flag was linearized by EcoR I digestion. The digested product was purified using a standard DNA purification kit, resolved on a 1% agarose gel, and quantified using a NanoDrop 2000 spectrophotometer by measuring OD and the OD /OD ratio.

*In vitro* transcription was performed using the iScribe™ T7 ARCA mRNA Kit (NEB, MA, USA), with N1-methylpseudouridine-5′-triphosphate (N1-mψUTP) substituting for UTP. mRNA concentration was determined using a Nanodrop2000 spectrophotometer, and transcript integrity was assessed by 5% urea denaturing PAGE.

#### Preparation of mRNA-LNPs

mRNA-LNPs were prepared by dissolving lipid01, DSPC, cholesterol, and DMG-PEG2k in anhydrous ethanol at a molar ratio of 20:40:38.5:1.5 to form the organic phase. The mRNA encoding the antigenic protein (OmpX-Fc mRNA) was diluted in sodium citrate buffer (50 mM, pH 4.0) to form the aqueous phase (N/P=6; charge molar ratio of mRNA to lipid01= 1.5:2) [16, 28–30]. The two phases were mixed at a 3:1 (v/v, aqueous:ethanol) ratio using a microfluidic device at a flow rate of 20 mL/min. The ethanol concentration was diluted 40-fold (to <1%) using sterile 1× D-PBS (without calcium and magnesium). The solution was transferred to a pre-sterilized Amicon^®^Ultra-15 centrifugal filter (MWCO 100 kDa) and centrifuged at 4,000×g for 15-30 min at 4°C. Sterile 1× PBS was added to achieve a final mRNA-LNP concentration of 2 mg/mL. Samples were stored at 4°C and used within one week.

#### Characterization of mRNA-LNP physicochemical properties

mRNA encapsulation efficiency was determined using the Quant-iT™ RiboGreen™ RNA Kit (Invitrogen, USA). This ultrasensitive fluorescent nucleic acid kit detects 1-200 ng of free nucleic acid without penetrating LNPs. Total nucleic acid content was measured by treating mRNA-LNPs with 1% Triton X-100 to release encapsulated RNA. Encapsulation efficiency (%) = [(nucleic acid after demulsification − nucleic acid before demulsification) / nucleic acid after demulsification] × 100.

For transmission electron microscopy (TEM), 500 μL of 4% phosphotungstic acid was mixed with an equal volume of mRNA-LNPs and sonicated for 5 min. A drop of the mixture was placed on a 200-mesh carbon-coated copper grid, and excess liquid was removed with filter paper. The grid was air-dried, rinsed with deionized water, air-dried again, and examined by TEM.

#### Cytotoxicity assay

HEK 293T cells were seeded in 96-well plates at 1×10□ cells/well and incubated for 12 h. Cells were then treated with 25 μg of mRNA-LNPs (SP-OmpX-Fc mRNA-LNPs) and incubated for 24 h at 37°C with 5% CO□. CCK-8 reagent (10 μL/well) was added, and cells were incubated for 1 h at 37°C with 5% CO□, after which OD□□□ was measured.

#### Western blotting

HEK 293T cells (3×10□) were seeded in 10 cm culture dishes and incubated at 37°C with 5% CO□. At 80-85% confluence, OmpX-Fc mRNA-LNPs (20 μg) were added dropwise, gently mixed, and incubated for 24 h at 37°C with 5% CO□. Untransfected HEK 293T cells served as negative controls.

Supernatants and cells were collected separately, and cells were lysed. OmpX-Fc protein was purified from both fractions using the FLAG Tag (DYKDDDDK) Fusion Protein Purification Kit.

Purified OmpX-Fc protein (20 μg) was mixed with 5× sampling buffer, denatured at 95°C for 10 min, separated by 10% SDS-PAGE, and transferred onto a 0.45 μm PVDF membrane by wet transfer. The membrane was blocked with 5% skim milk for 1 h and washed three times with PBST (containing 0.1% Tween-20). The membrane was then incubated with an anti-Flag mouse monoclonal antibody, followed by goat anti-mouse IgG H&L (DyLight^®^ 488) secondary antibody (1:4,000 dilution), with gentle shaking at room temperature (protected from light) for 1 h, and washed three times with TBST. Finally, the membrane was imaged.

#### Indirect immunofluorescence

The OmpX-Fc recombinant plasmid was purified using an endotoxin-free plasmid extraction kit (Fast High Purity) and adjusted to 1,000 μg/mL in TE buffer. OmpX-Fc mRNA-LNPs (2.5 μg mRNA) were transfected into HEK 293T cells for 24 h, and protein expression was detected by immunofluorescence staining. Transfected cells were fixed with 4% formaldehyde at room temperature for 10 min and washed twice with PBST. Cells were incubated with 2% BSA and 0.3% Triton X-100 for 30 min at room temperature, washed twice with PBST, and incubated with anti-Flag mouse monoclonal antibody at 4°C overnight. Cells were then incubated with goat anti-mouse IgG H&L (DyLight^®^ 488) secondary antibody (1:4,000 in PBST) at room temperature (protected from light) for 1 h. After washing with PBST, cells were mounted with anti-fluorescence-attenuating mounting medium containing DAPI, and fluorescence signals were visualized by confocal microscopy.

#### Immunoassay

Specific pathogen-free, 6-8-week-old female C57BL/6 mice were purchased from Spearfish Biotechnology Co. (Beijing, China). Mice were housed in a controlled environment (relative humidity 55 ± 5%, 22°C, 12 h:12 h light/dark cycle) with ad libitum access to food and water. Mice were randomly divided into four groups (n = 17): alum (0.36%), luciferase mRNA-LNPs (5 μg mRNA), OmpX-Fc mRNA-LNPs (5 μg mRNA), and OmpX protein (80 μg). Vaccines were administered by intramuscular injection, and mice were immunized three times at 14-day intervals.

#### Indirect ELISA for serum antibody titers

Levels of OmpX-specific IgG, IgG1, and IgG2a antibodies in serum samples collected on days 14, 28, and 42 after immunization were measured by indirect ELISA. OmpX-Fc or OmpX protein (500 ng/well) was coated onto Corning™ Costar™ 9018 ELISA plates using coating buffer, and plates were incubated at 4°C overnight. Wells were washed three times with 0.05% PBST, blocked with 3% BSA-PBST (100 μL/well) for 1 h at 37°C, and washed again. Mouse serum diluted 1:100 in PBS (100 μL/well) was added and incubated at 37°C for 1 h. After washing, HRP-conjugated goat anti-mouse IgG (H+L) secondary antibody (1:4,000) was added (100 μL/well) and incubated at 37°C for 1 h. After washing, 100 μL of TMB substrate solution was added and allowed to react for 15 min at 37°C. The reaction was stopped with 50 μL of 2 M H□SO□, and OD□□□ was measured.

For IgG1 and IgG2a detection, the secondary antibodies were replaced with HRP-conjugated goat anti-mouse IgG1 (1:2,000) and HRP-conjugated goat anti-mouse IgG2a (1:500), respectively. All ELISAs were coated with tag-free recombinant OmpX (500 ng/well). To exclude reactivity against the human IgG1 Fc tag present in the immunogen, a parallel plate coated with human IgG1 Fc (500 ng/well) was included, and Fc-reactive background was subtracted; sera with Fc-only reactivity >10% of the OmpX reactivity were flagged.

#### Cytokine detection in serum

Levels of IL-4, IFN-γ, and IL-17A in serum samples collected on days 14, 28, and 42 were measured using commercial ELISA kits (Elabscience, China) according to the manufacturer’s instructions.

#### Splenic lymphocyte proliferation assay

Splenocyte proliferation was assessed using a CCK-8 kit according to the manufacturer’s protocol. Splenic lymphocytes were isolated using a mouse lymphocyte isolation medium kit (Solarbio). Cells were plated in 96-well plates (100 μL/well) at a density of 5×10□ cells/well and stimulated with purified OmpX-Fc or OmpX protein (2 μg/mL). The negative control received 100 μL of DMEM without treatment; the positive control consisted of normal mouse splenocytes treated with concanavalin A (ConA, 2 μg/mL, MCE). Cells were cultured at 37°C with 5% CO□ for 72 h. The medium was then replaced with fresh medium, CCK-8 solution (10 μL/well) was added, and plates were incubated for 4 h. Absorbance was measured at 450 nm using a microplate reader. All treatments were performed in triplicate. Cytokines (IL-4, IFN-γ, and IL-17A) in the collected culture supernatant were measured by indirect ELISA.

#### Flow cytometric immunophenotyping

Flow cytometry was used to analyze the percentages of CD3□ CD4□ T and CD3□ CD8a□ T lymphocytes in mouse spleens. Lymphocytes were isolated using a mouse lymphocyte isolation medium kit (Solarbio). Cells were incubated with FITC anti-mouse CD3 (0.5 μg/10□ cells), APC anti-mouse CD4 (0.25 μg/10□ cells), and PerCP anti-mouse CD8a (0.25 μg/10□ cells) antibodies for 30 min at 4°C in the dark. Labeled cells were washed three times with pre-cooled PBS and analyzed by flow cytometry.

#### Intracellular cytokine staining

Mice were euthanized 14 days after the third immunization, and spleens were collected immediately. Single-cell suspensions were prepared, washed with PBS, and lymphocytes were isolated using the mouse lymphocyte isolation medium kit. Cells were counted and seeded at 2×10 cells/well in 6-well plates. OmpX-Fc or OmpX protein (2 μg/mL) was added for 6 h of stimulation, followed by brefeldin A for an additional 6 h. Cells were washed with PBS containing 0.2% BSA, resuspended in 100 μL, and incubated with anti-CD16/CD32 antibodies for 30 min. After washing, cells were incubated with FITC anti-mouse CD3 (0.5 μg/10□ cells), APC anti-mouse CD4 (0.25 μg/10 cells), and PerCP anti-mouse CD8a (0.25 μg/10□ cells) at 4°C for 30 min in the dark. Intracellular cytokine staining was performed using a Cytofix/Cytoperm Fixation/Permeabilization Kit (BD Biosciences, USA). Cells were then incubated with PE/Cyanine7 anti-mouse IL-4 (1.0 μg/10□ cells), PE/Cyanine7 anti-mouse IFN-γ (1.0 μg/10□ cells), or PE/Cyanine7 anti-mouse IL-17A (0.25 μg/10□ cells) for 30 min at 4°C in the dark. After centrifugation, cells were washed with pre-cooled PBS and analyzed by flow cytometry. Data were analyzed using FlowJo v10.7.1.

#### Challenge infection experiments

Bacterial challenge was performed two weeks after the third immunization. For the sepsis model, each mouse received an intraperitoneal injection of 5×10^8^ CFU of KP suspension and was observed for 14 days.

#### Bacterial load in tissues

Mice were euthanized 24 h after KP challenge, and venous blood was collected. The liver, lung, and spleen were harvested, weighed, and homogenized in 1 mL of sterile PBS. Tissue homogenates were serially diluted, plated on LB agar, and incubated at 37°C overnight. Bacterial colonies (CFU) were counted the following day.

### *In vitro* bacterial inhibition tests

The protocol was adapted as follows: sera from immunized mice were heat-inactivated at 56°C for 1 h to remove endogenous complement activity. KP (10□ CFU) was incubated with 50 μL of a 1:100 dilution of heat-inactivated mouse serum for 30 min at 37°C, after which baby rabbit complement (e.g., 10% v/v; Cedarlane) was added and the mixture was incubated for a further 30, 60, or 90 min. Reactions were terminated by adding 950 μL of LB medium; the mixture was spread onto LB plates and incubated at 37°C overnight. Controls consisted of (i) bacteria incubated without serum, (ii) bacteria incubated with complement alone (no serum), and (iii) bacteria incubated with serum plus 10 mM EDTA to block complement activation. Percentage of lysis = 100 − 100 × [(CFU of experimental wells) / (CFU of control wells)].

### Histological analysis

Mice were euthanized by isoflurane inhalation, and major organs (heart, liver, spleen, lung, and kidney) from immunized and control (alum and luciferase mRNA-LNP) groups were collected in 4% paraformaldehyde for paraffin sectioning and hematoxylin and eosin (H&E) staining.

### Statistical analysis

Statistical analysis was performed using Prism 8.0 (GraphPad Software). Multiple group comparisons were performed using one-way ANOVA followed by Tukey’s multiple comparison post hoc test. Between-group comparisons were performed using unpaired Student’s t-test. *P* < 0.05 was considered statistically significant (\**P* < 0.05, \*\**P* < 0.01, \*\*\**P* < 0.001). NS, not significant. **RESULTS**

### OmpX vaccine design and conservation analysis across KP subtypes

The OmpX mRNA vaccine was constructed with the SP-OmpX-Fc-Flag expression cassette under the control of the T7 promoter, incorporating a 5’ cap, 5’ UTR, signal peptide sequence, OmpX antigen-encoding region, human Fc tag, 3’ UTR, and polyA tail (Fig. 1A). Bioinformatics analysis of OmpX sequences from a comprehensive panel of KP clinical strains revealed that OmpX is highly conserved across all three major clinical subtypes-classical (cKP), hypervirulent (hvKP), and multidrug-resistant (MDR) strains-with greater than 95% amino acid sequence identity among isolates (Fig. 1B). This broad conservation suggests that an OmpX-based vaccine could provide serotype-independent protection across diverse KP lineages.

**Fig. 1.**
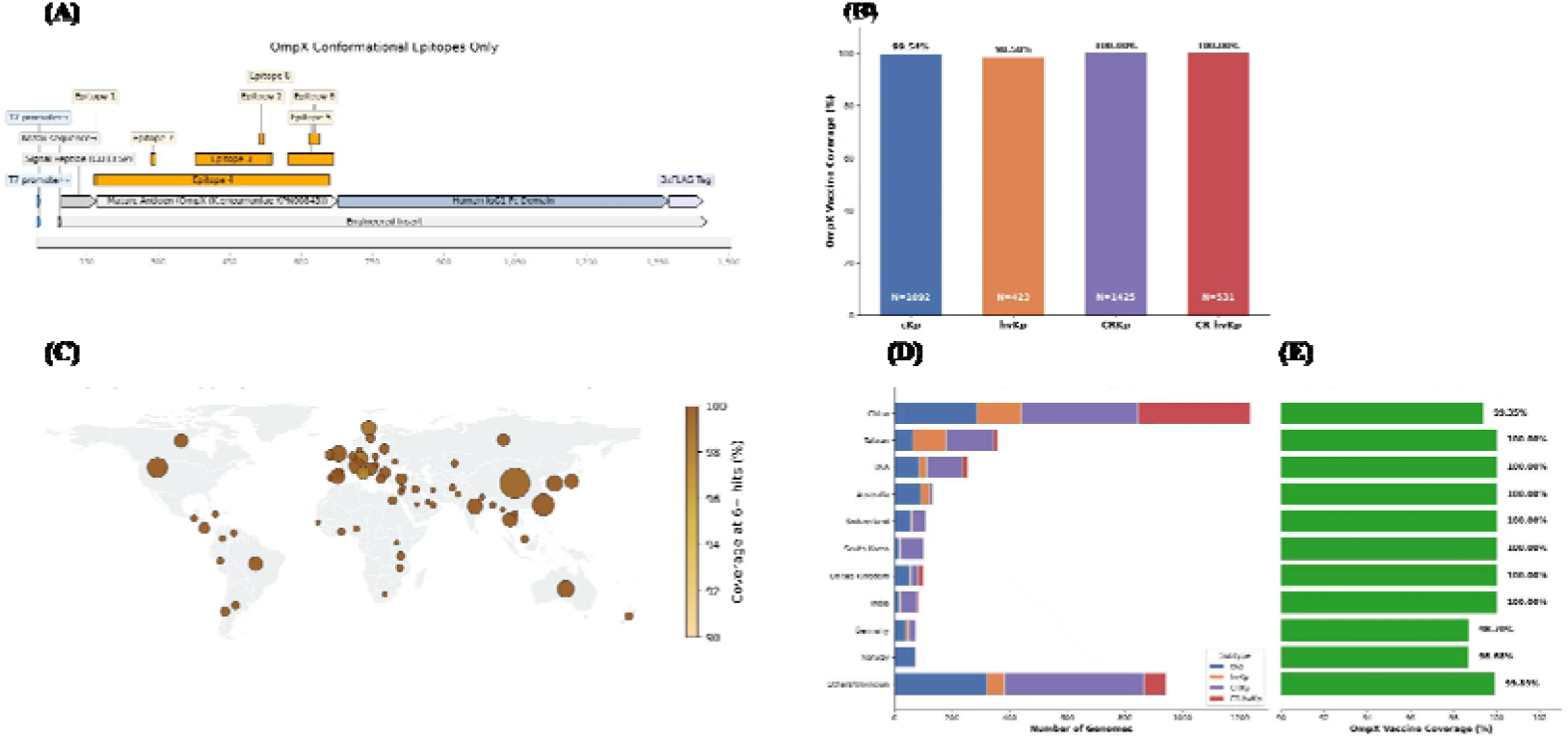
Schematic diagram of OmpX vaccine construction and the conserved types and regional distribution of OmpX epitopes. (A) Schematic diagram of the OmpX vaccine plasmid construction; (B) universal coverage of the OmpX mRNA-LNP vaccine across clinical subtypes; (C) geographical mapping of clinical strains and vaccine coverage; (D) KP clinical subtype distribution by region; (E) conservation of OmpX vaccine epitope regions.

Geographic mapping of KP clinical strains demonstrated that OmpX is ubiquitously distributed across Asia, Europe, North America, Africa, and Oceania, with robust predicted vaccine coverage in all regions (Fig. 1C). Subtype distribution analysis revealed regional variation in the relative prevalence of cKP, hvKP, and MDR strains; MDR strains were most prevalent in Asia, whereas hvKP strains were enriched in the Asia-Pacific region (Fig. 1D). Importantly, the predicted B-cell and T-cell epitope regions of OmpX were highly conserved across all geographic isolates, with less than 2% amino acid variation in epitope-containing sequences (Fig. 1E). These results indicate that OmpX is a conserved, broadly distributed antigen suitable for development as a universal KP vaccine target.

### Immunogenicity, antigenicity, allergenicity, and toxicity of OmpX

The IEDB immunogenicity server predicted an immunogenicity score of 0.86 for the OmpX vaccine, and the ANTIGENpro server predicted an antigenicity score of 1.01. AllerTOP v.2.0, AllergenFP v.1.0, and ToxinPred predicted that OmpX is non-sensitizing and non-toxic. These results indicate that OmpX is a promising KP vaccine candidate with high antigenicity, strong immunogenicity, and favorable safety characteristics.

### Physicochemical properties and solubility

Physicochemical parameters predicted by the ExPASy ProtParam server are summarized in Table 1. The predicted solubility score of the OmpX vaccine was 0.654 (Fig. 2A), indicating good solubility.

**Fig. 2.**
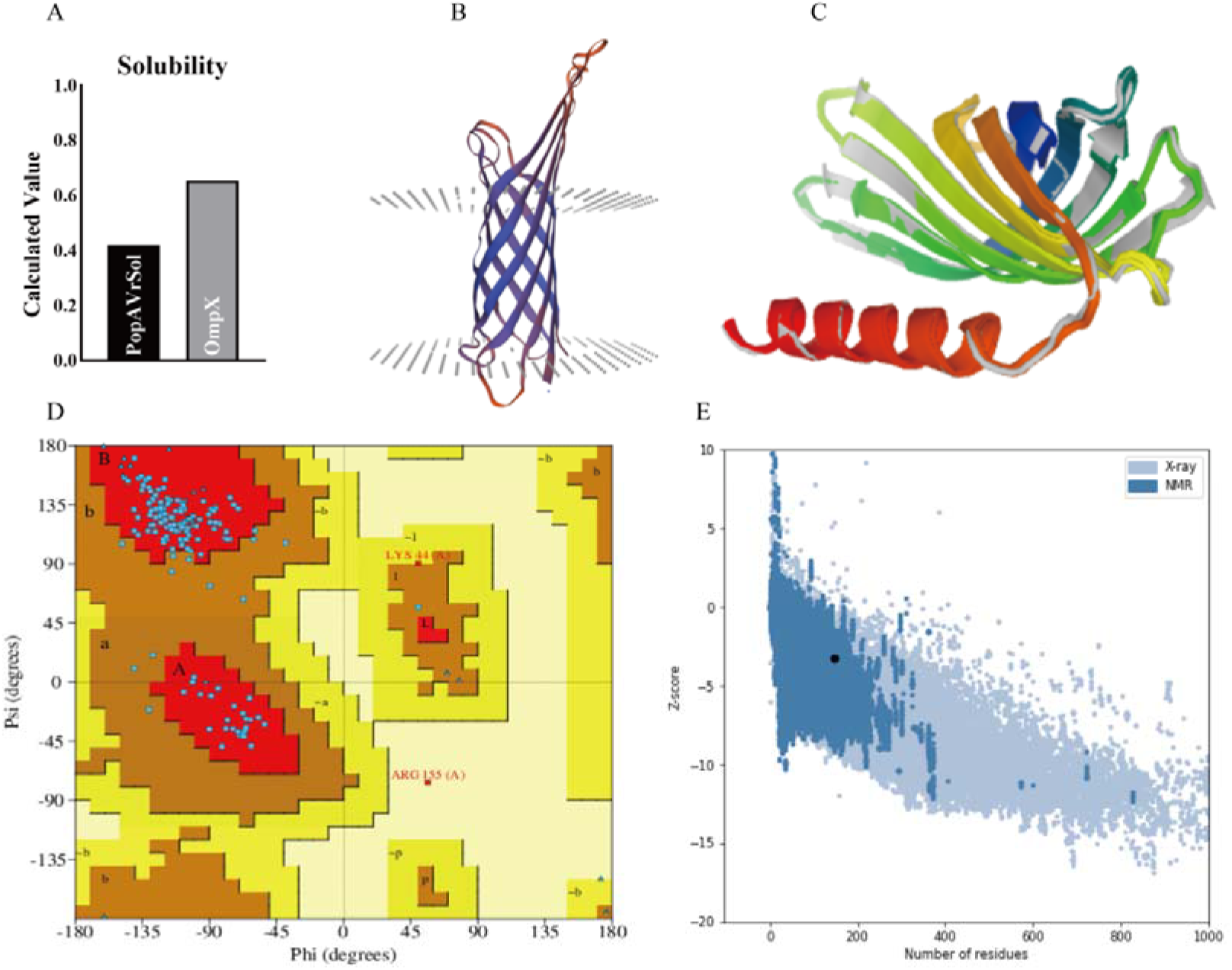
Solubility and tertiary structure of the OmpX vaccine. (A) Solubility of the OmpX vaccine predicted by the Protein-Sol server; (B) tertiary structure of the OmpX vaccine predicted by the 3Dpro server before optimization; (C) optimized tertiary structure of the OmpX vaccine predicted by the GalaxyRefine web server; (D) Ramachandran plot analysis of the OmpX vaccine generated with PROCHECK; (E) ProSA web server analysis of the Z-score of the OmpX vaccine.

**Table 1.**
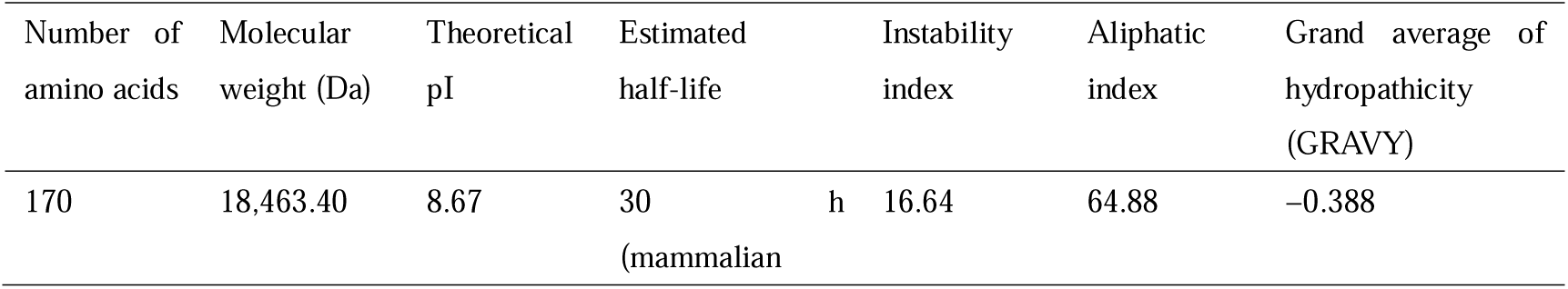
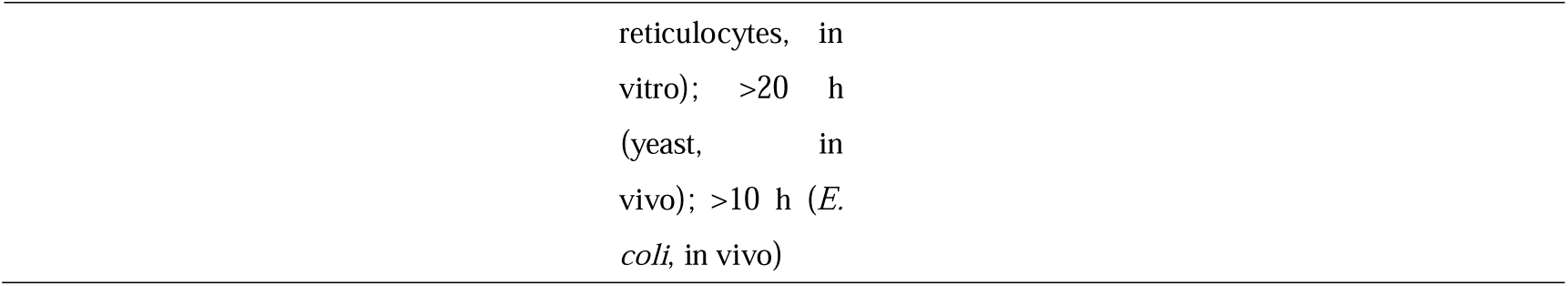
Physicochemical parameters of the OmpX vaccine predicted by the ExPASy ProtParam server.

### Validation of the predicted secondary and tertiary structures of OmpX

The secondary structure of OmpX was composed of 56.32% α-helices, 10.6% β-sheets, and 33.08% random coils. The initial 3D model was generated using the 3Dpro server (Fig. 2B) and subsequently refined using the GalaxyRefine web server. Refined models were ranked by GDT-HA and MolProbity scores (Table 2). Model quality was evaluated using five parameters-GDT-HA (backbone accuracy), RMSD (structural deviation relative to a reference), clash score (clashes per 1,000 atoms), percentage of poor rotamers, and Ramachandran plot quality-with higher GDT-HA and lower MolProbity values indicating superior model quality (Fig. 2C), which is critical for downstream analyses.

**Table 2.** Model parameters for GalaxyRefine web server refinement.

| Model | GDT-HA | RMSD | MolProbity | Clash score | Poor rotamers (%) | Ramachandran favored (%) |
| --- | --- | --- | --- | --- | --- | --- |
| Initial | 1.0000 | 0.000 | 1.212 | 1.6 | 1.5 | 97.0 |
| Model 1 | 0.9544 | 0.430 | 1.673 | 7.3 | 2.3 | 98.8 |
| Model 2 | 0.9632 | 0.393 | 1.688 | 5.7 | 1.5 | 96.4 |
| Model 3 | 0.9574 | 0.409 | 1.439 | 6.5 | 0.8 | 97.6 |
| Model 4 | 0.9559 | 0.411 | 1.577 | 8.0 | 1.5 | 98.2 |
| Model 5 | 0.9485 | 0.422 | 1.692 | 8.8 | 1.5 | 97.6 |

### Validation of the tertiary structure and molecular docking

The overall quality factor of the refined OmpX model, as assessed by ERRAT, was 85.36%. Ramachandran plot analysis showed that 88.7% of residues were in the most favored region, 9.9% in additionally allowed regions, and 0.7% in generously allowed regions, with an overall quality factor of 98.6% (Fig. 2D). The OmpX model had a Z-score of −3.26 (Fig. 2E). Molecular docking analysis demonstrated stable binding between OmpX and HLA-DRB1*01:01 (1AQD), TLR-2 (6NIG), and TLR-4 (4G8A) (Fig. 3A-C).

**Fig. 3.**
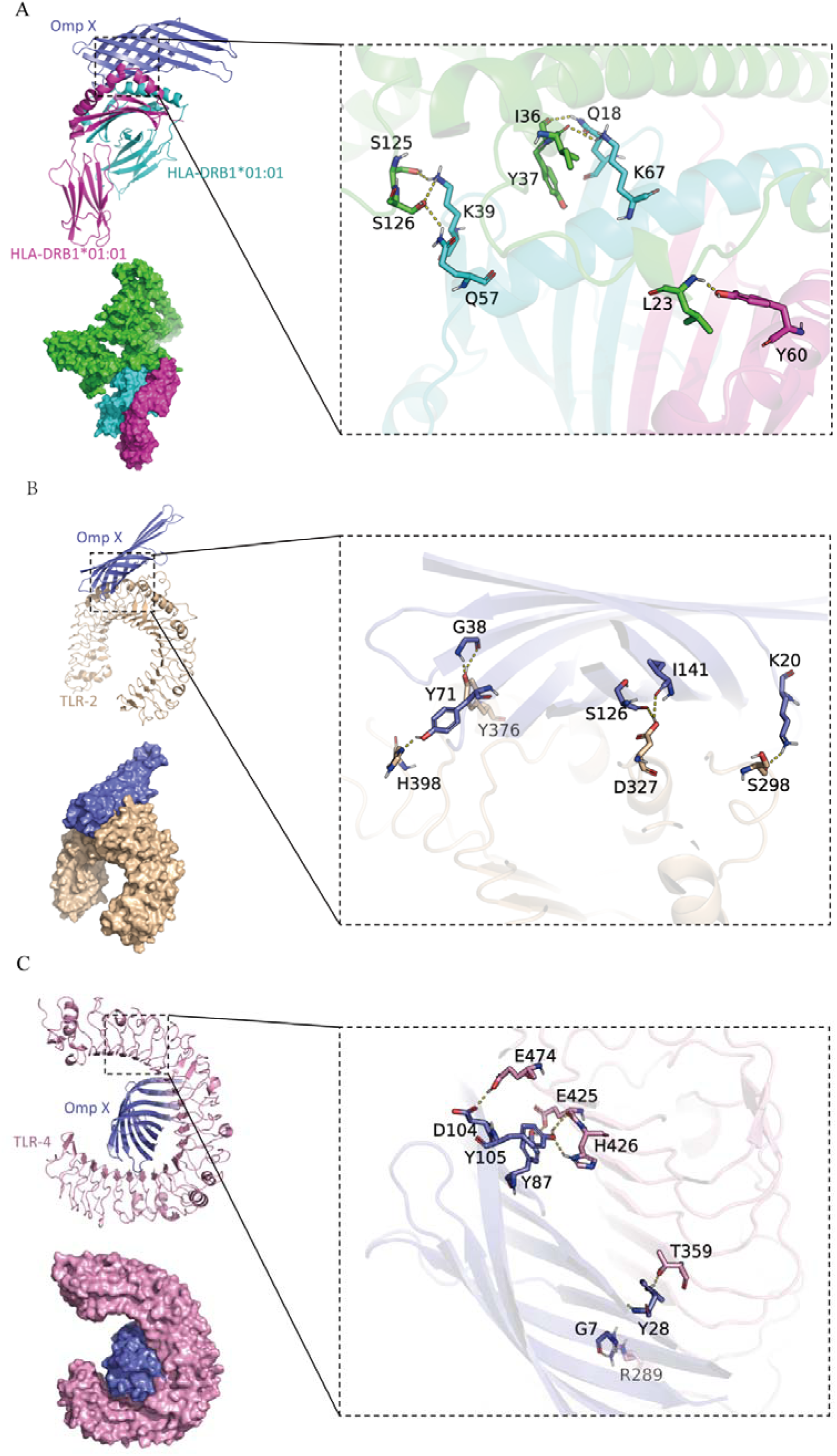
Molecular docking between the OmpX vaccine and HLA-DRB1*01:01, TLR-2, and TLR-4. (A) Molecular docking between the OmpX vaccine and HLA-DRB1*01:01; (B) molecular docking between the OmpX vaccine and TLR-2; (C) molecular docking between the OmpX vaccine and TLR-4.

### Conformational B-cell epitopes

The OmpX vaccine formed eight conformational B-cell epitopes with scores ranging from 0.557 to 0.975 (Fig. 4), encompassing 79 residues in total (Table 3). The three highest-scoring epitopes comprised residues M1-K3 (score 0.975), Y118-H120 (0.963), and D73-D123 (0.776). The predicted epitopes cover 79/170 (46.5%) of the OmpX sequence.

**Fig. 4.**
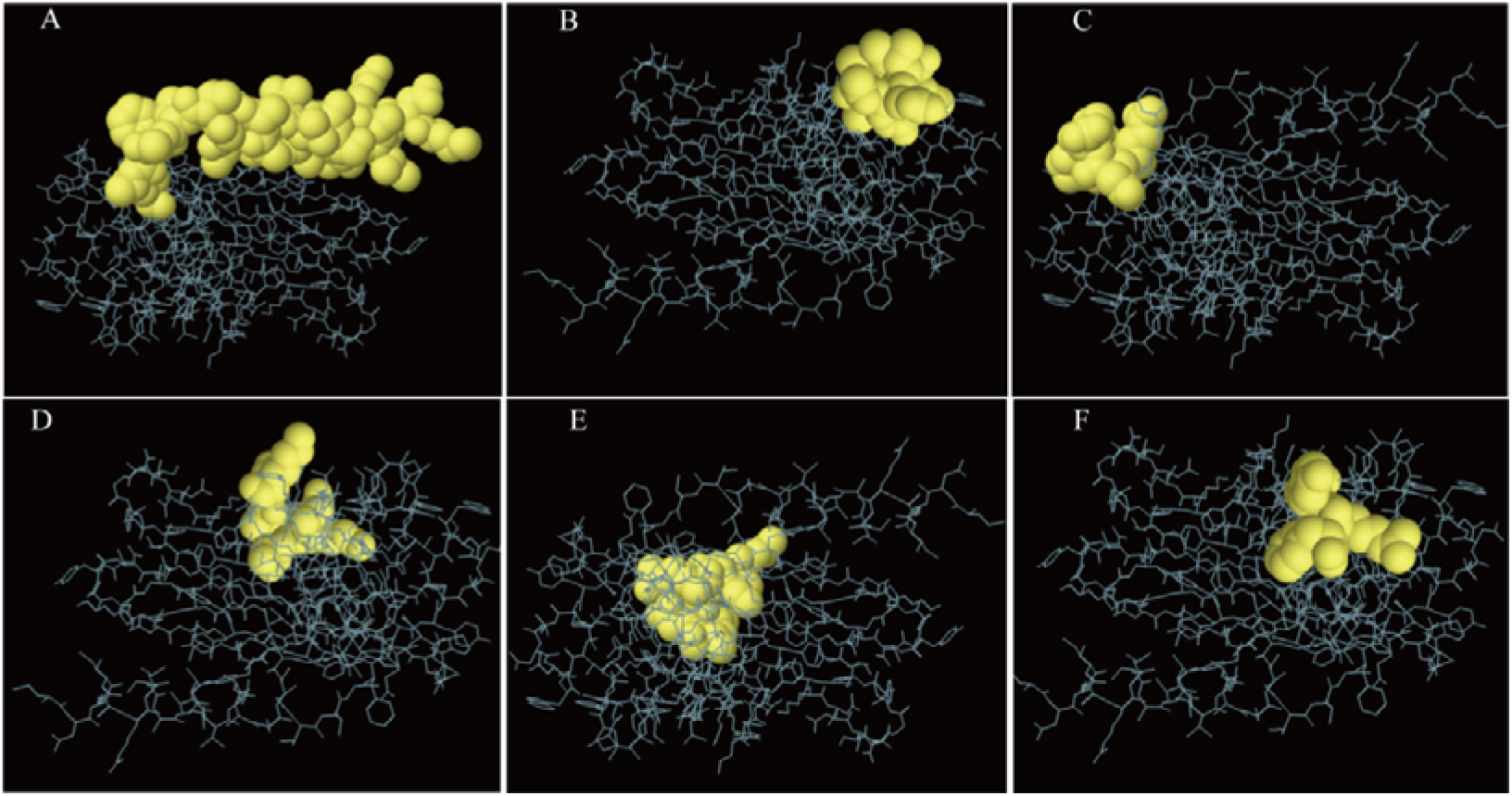
Conformational B-cell epitopes predicted by the ElliPro server. (A-F) Locations of the OmpX conformational B-cell epitopes; the remaining regions are shown in gray.

**Table 3.**
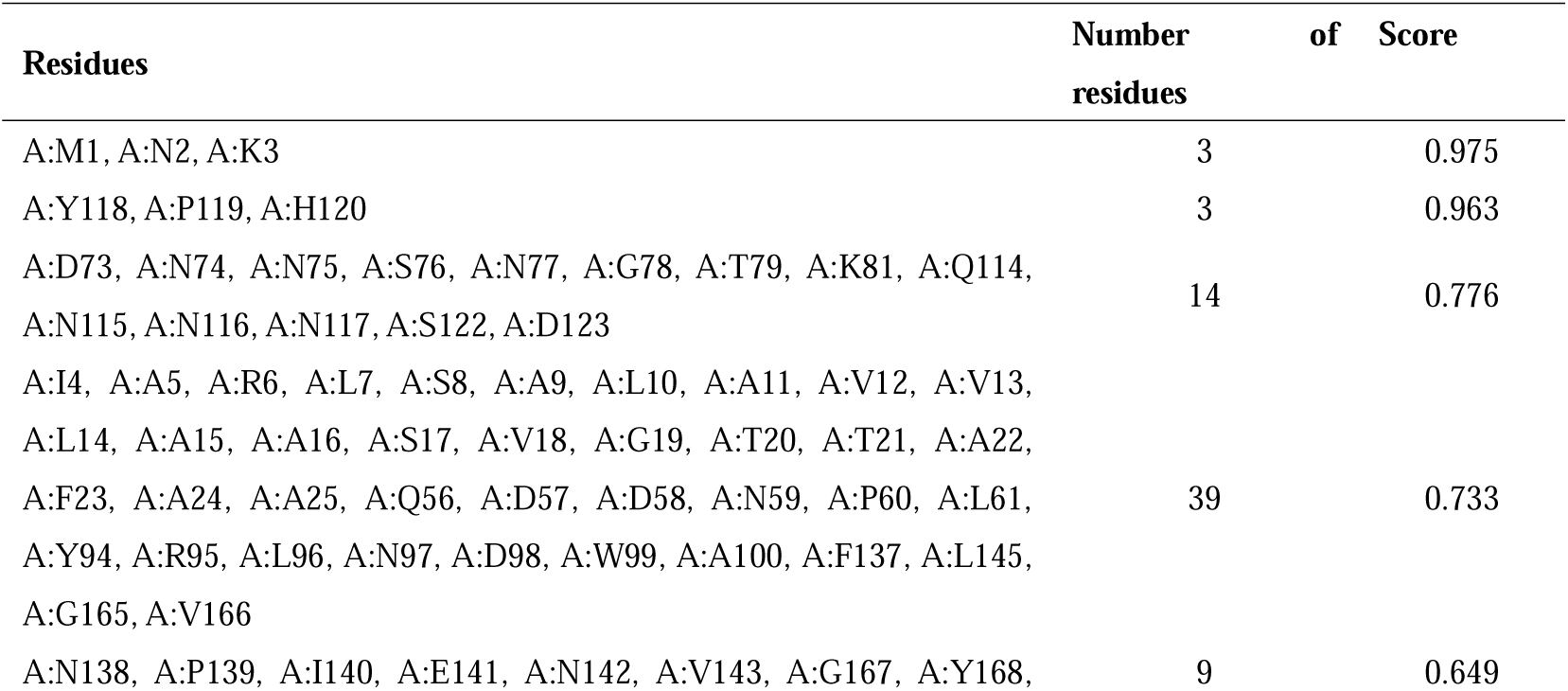
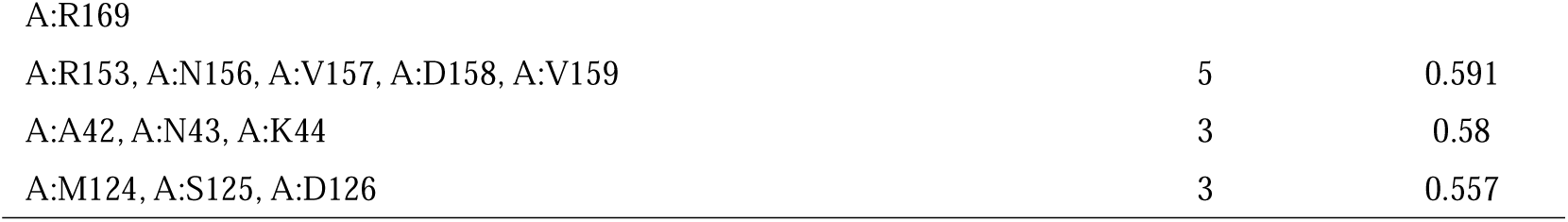
ElliPro-predicted conformational B-cell epitopes of the OmpX vaccine.

The OmpX vaccine significantly activated dendritic cells (DCs)-the most potent antigen-presenting cells (APCs)-which are essential for presenting vaccine antigens to T cells. The total DC count in the OmpX vaccine group remained at approximately 200 cells/mm^3^ after stimulation (Fig. 5A). Macrophages were also significantly activated from the first to the third injection, with active macrophage counts remaining at approximately 195 cells/mm^3^ (Fig. 5B).

**Fig. 5.**
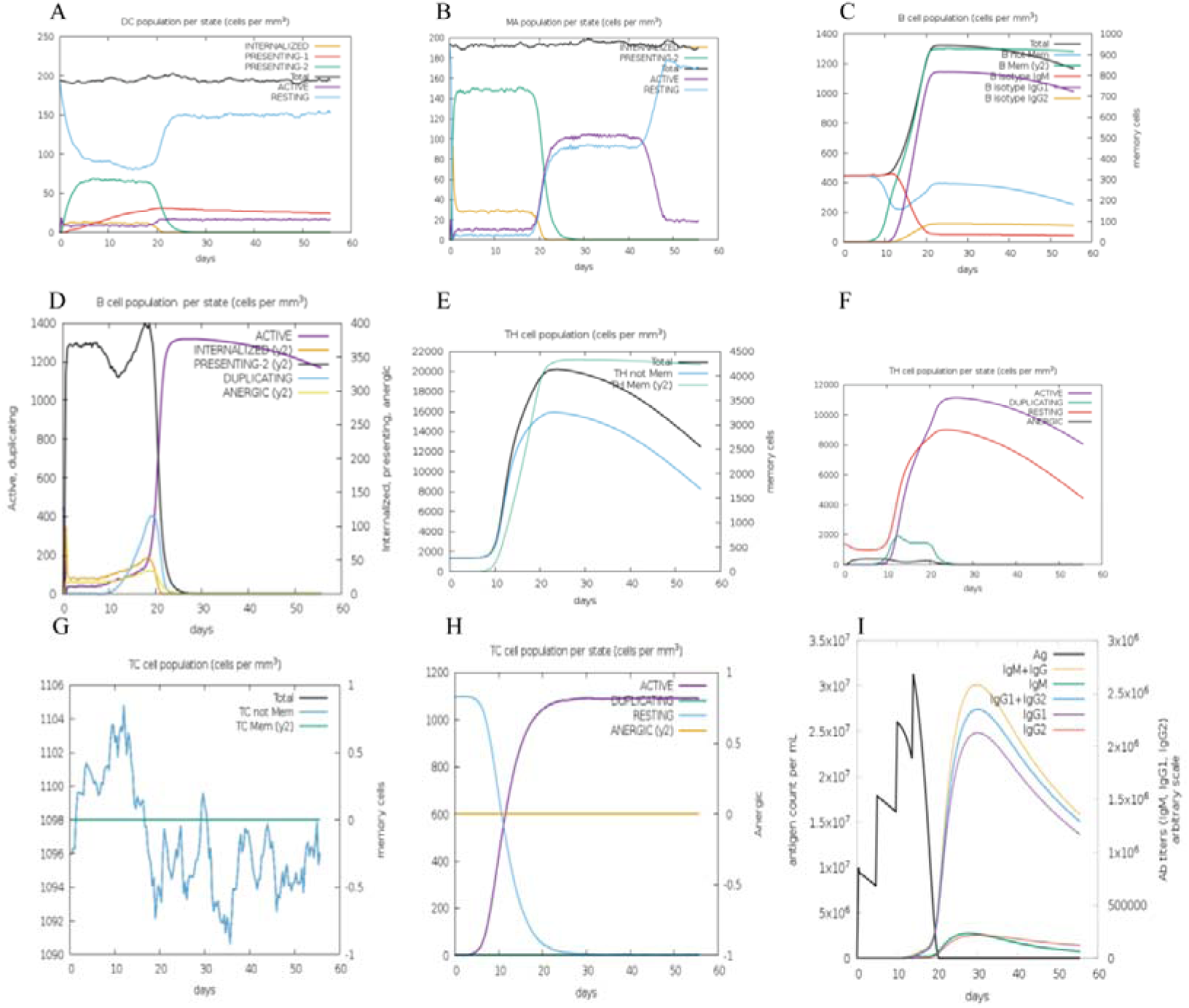
Immune simulation of innate immune cells predicted by the C-ImmSim server. (A) Number and status of DCs after OmpX vaccine immunization; (B) number and status of macrophages after OmpX vaccine immunization; (C) B-cell populations after OmpX vaccine immunization; (D) Th-cell populations after OmpX vaccine immunization; (E) status of Th-cell populations after OmpX vaccine immunization; (F) TC-cell populations after OmpX vaccine immunization; (G) status of TC-cell populations after OmpX vaccine immunization; (H) levels of OmpX-specific IgM; (I) levels of OmpX-specific IgG, IgG1, and IgG2 antibodies.

B cell counts in each activation state peaked at approximately 1,350 cells/mm^3^ after the third stimulation (Fig. 5C). Adaptive immune cells, including B cells, CD4□ T cells, and CD8□ T cells, play essential roles in resistance to KP. Although B cells are traditionally considered to play a secondary role in KP clearance, the humoral immunity they generate is nonetheless significant. The OmpX vaccine stimulated the formation of three prominent peaks of both non-memory and memory T helper (Th) cells, with peak counts of approximately 16,000, 21,000, and 20,000 cells/mm^3^ for non-memory Th cells, memory Th cells, and total Th cells, respectively, after the third stimulation (Fig. 5D). Active and resting CD4□ T cells peaked at approximately 11,000 and 9,000 cells/mm^3^, respectively, after the third stimulation (Fig. 5E).

CD8□ T cells contribute to KP clearance by producing cytotoxic factors such as perforin and granzyme B. Non-memory cytotoxic T lymphocytes (CTLs) reached a peak of approximately 1,106 cells/mm^3^ after the third stimulation (Fig. 5F). Active CTLs peaked after the third immunization, reaching approximately 1,100 cells/mm^3^ (Fig. 5G, H).

### Simulated humoral immune response to OmpX immunization

Simulated OmpX immunization produced markedly elevated levels of OmpX-specific antibodies. After the third injection, IgM levels peaked at approximately 2.5×10□ /mL and then gradually declined (Fig. 5H). Combined IgG1+IgG2 levels peaked at approximately 2.7×10□ /mL, followed by a gradual decrease. Individual IgG1 and IgG2 levels reached approximately 2.5×10□ /mL and 2.5×10 /mL, respectively (Fig. 5I).

### Physicochemical properties of the mRNA-LNPs

The mRNA vaccine encoding OmpX was modified to include a Cap1 (N7mGpppAm) sequence, a human signal peptide, and the Fc sequence (Fig. 6A). mRNA was synthesized by T7 polymerase-mediated in vitro transcription, substituting uridine (U) with N1-methylpseudouridine (m1ψ), and LNPs were prepared using microfluidics. The LNPs comprised four components-lipid01(Fig. S1, Fig. S2), DSPC, cholesterol, and DMG-PEG2k-at a molar ratio of 20:40:38.5:1.5. The mean particle size was 93.9 nm (Fig. 6B), with a polydispersity index of approximately 0.13 and an encapsulation efficiency of approximately 90% (Fig. 6D). The average surface charge was −4.815 mV at physiological pH, and CCK-8 assays confirmed low cytotoxicity (Fig. 6E). TEM revealed a homogeneous, solid, spherical morphology (Fig. 6C).

**Fig. 6.**
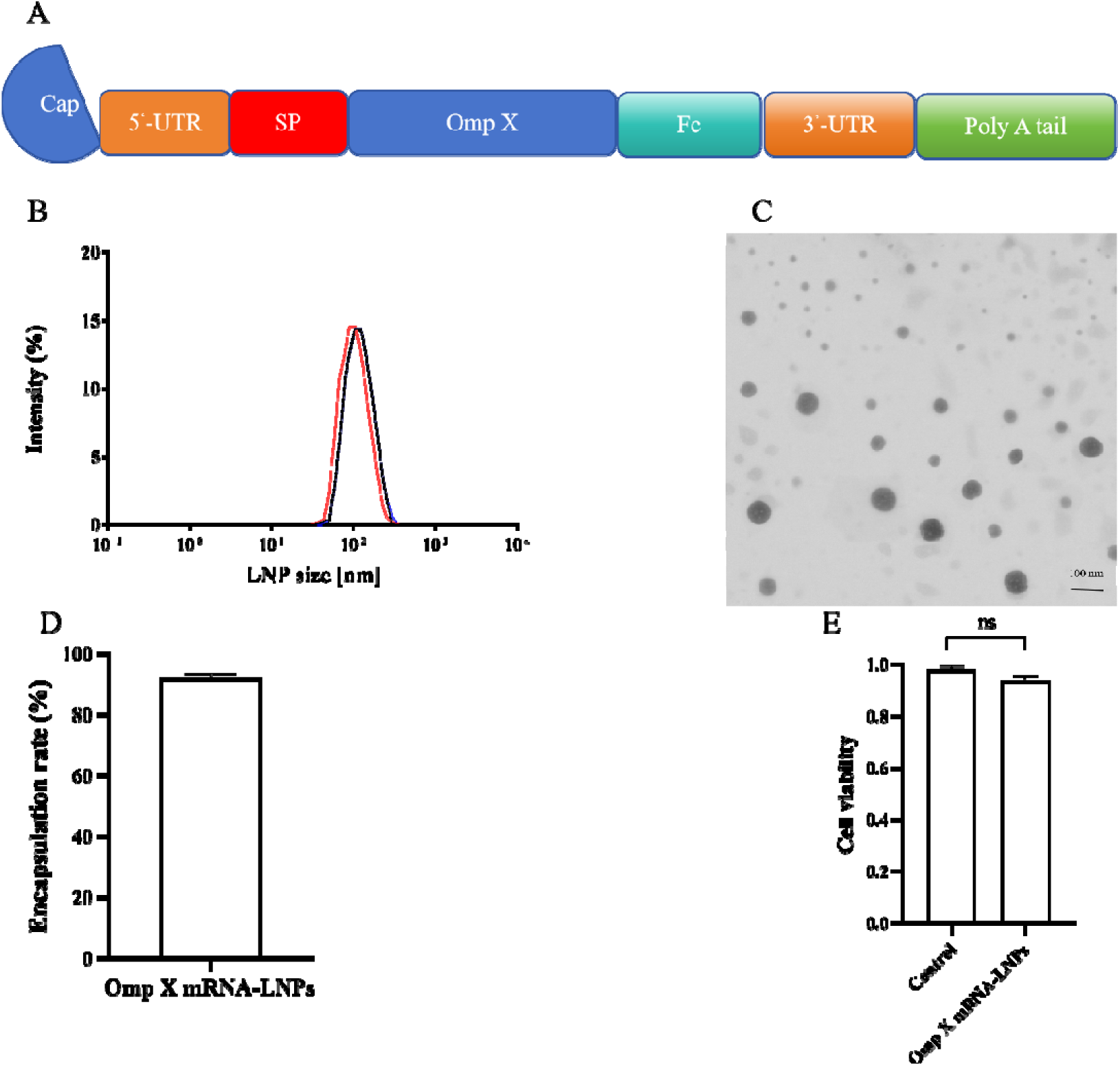
mRNA-LNP vaccine design and in vitro characterization. (A) Schematic illustration of the SP-OmpX-Fc mRNA construct. The mRNA construct consists of a 5’ cap followed by the 5’ UTR, signal peptide, OmpX coding sequence, 3’ UTR, and polyA tail. (B) Particle size of OmpX mRNA-LNPs. (C) Transmission electron microscopy (TEM) image of LNP-encapsulated OmpX-Fc mRNA (scale bar, 100 nm). (D) Encapsulation efficiency of mRNA-LNPs determined by the RiboGreen assay. (E) Cytotoxicity of blank LNPs and mRNA-LNP vaccines in HEK 293T cells. Data are presented as means ± SEM. \**P*<0.05; \*\**P*<0.01; \*\*\**P*<0.001; \*\*\*\**P*<0.0001; ns, not significant.

### Expression of OmpX-Fc mRNA-LNPs in HEK 293T cells

HEK 293T cells transfected with OmpX-Fc mRNA-LNPs showed green fluorescence, whereas control cells showed no visible fluorescence (Fig. 7A). Transfection with OmpX-Fc mRNA-LNPs resulted in significantly higher OmpX-Fc protein levels (Fig. 7B, C), confirming efficient protein expression.

**Fig. 7.**
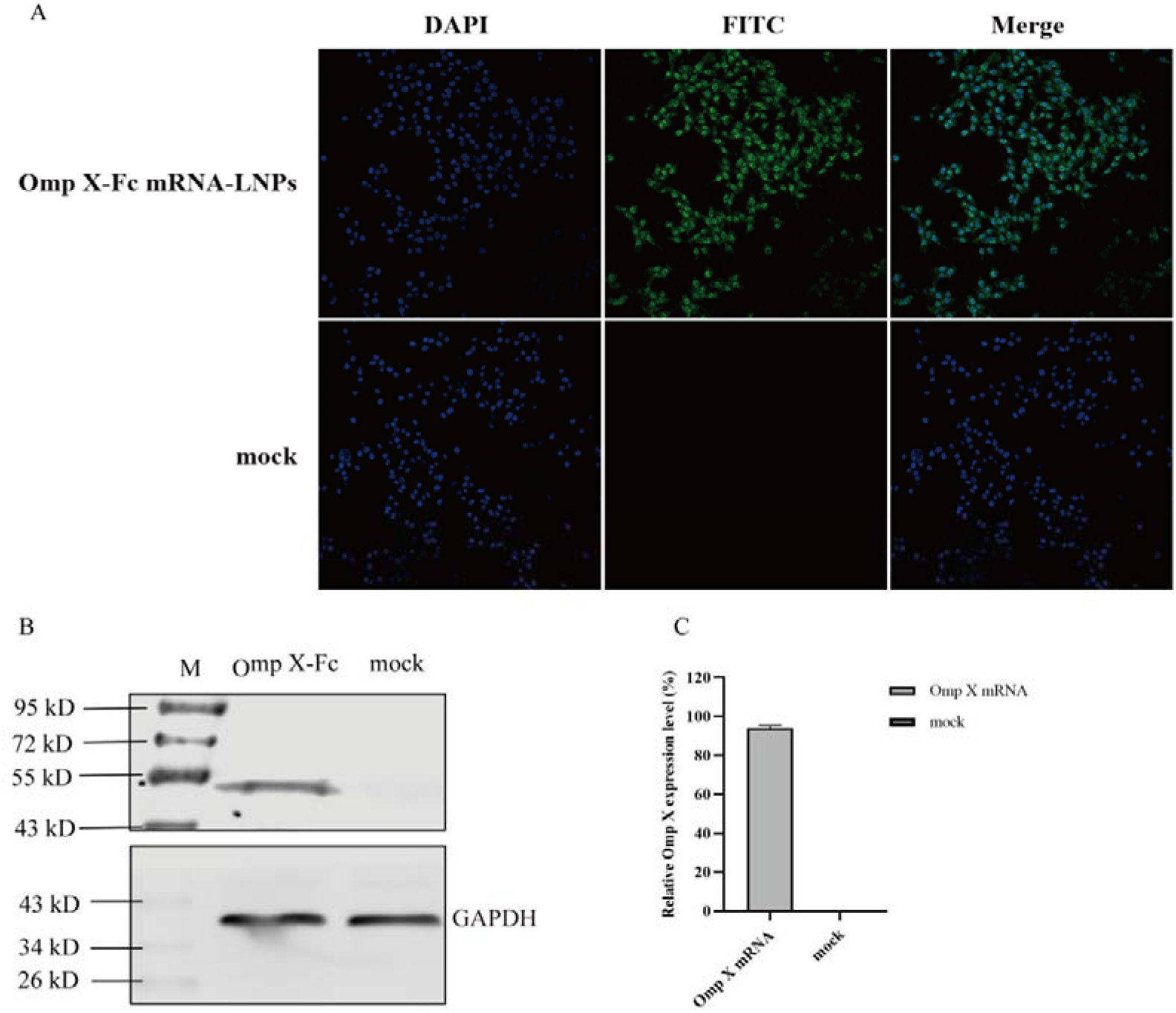
*In vitro* expression of mRNA-LNPs. (A) HEK 293T cells were treated with mRNA-containing LNPs for 24 h and then fixed for immunofluorescence assay (IFA) or lysed for protein blotting. (B) Anti-Flag antibody was used as the primary antibody for IFA and immunoblotting; the secondary antibody was labeled with green fluorescence, and nuclei were stained with DAPI (blue). (C) Quantification of OmpX-Fc expression.

### Immunogenicity and humoral immune response induced by OmpX-Fc mRNA-LNPs

Previous studies have shown that heat-killed KP can induce lung-resident memory (TRM) cells, a subset of which recognizes bacterially encoded outer membrane proteins. We therefore investigated whether this approach could be used to develop an mRNA vaccine against KP infection. OmpX was cloned from a KP strain, and immunized mice were challenged to assess serotype-independent immunity.

The immunization schedule is shown in Fig. 8A. Neither OmpX-Fc mRNA-LNP nor OmpX protein vaccination caused observable adverse effects on mouse appearance or behavior. By day 14, both the OmpX-Fc mRNA-LNP and OmpX protein groups had detectable serum IgG against OmpX. IgG titers increased after the first immunization, peaked on day 28 and remained elevated through day 42 in the immunized groups, while remaining undetectable in the alum and luciferase mRNA-LNP control groups (Fig. 8B-D). Antibody levels were significantly higher in both vaccine groups than in controls (*P* < 0.001). These results confirm that both OmpX-Fc mRNA-LNPs and OmpX protein are immunogenic and capable of inducing OmpX-specific IgG in mice. Further evaluation showed significant increases in CD4□ and CD8a□ T cells in the mRNA vaccine group compared with controls.

**Fig. 8.**
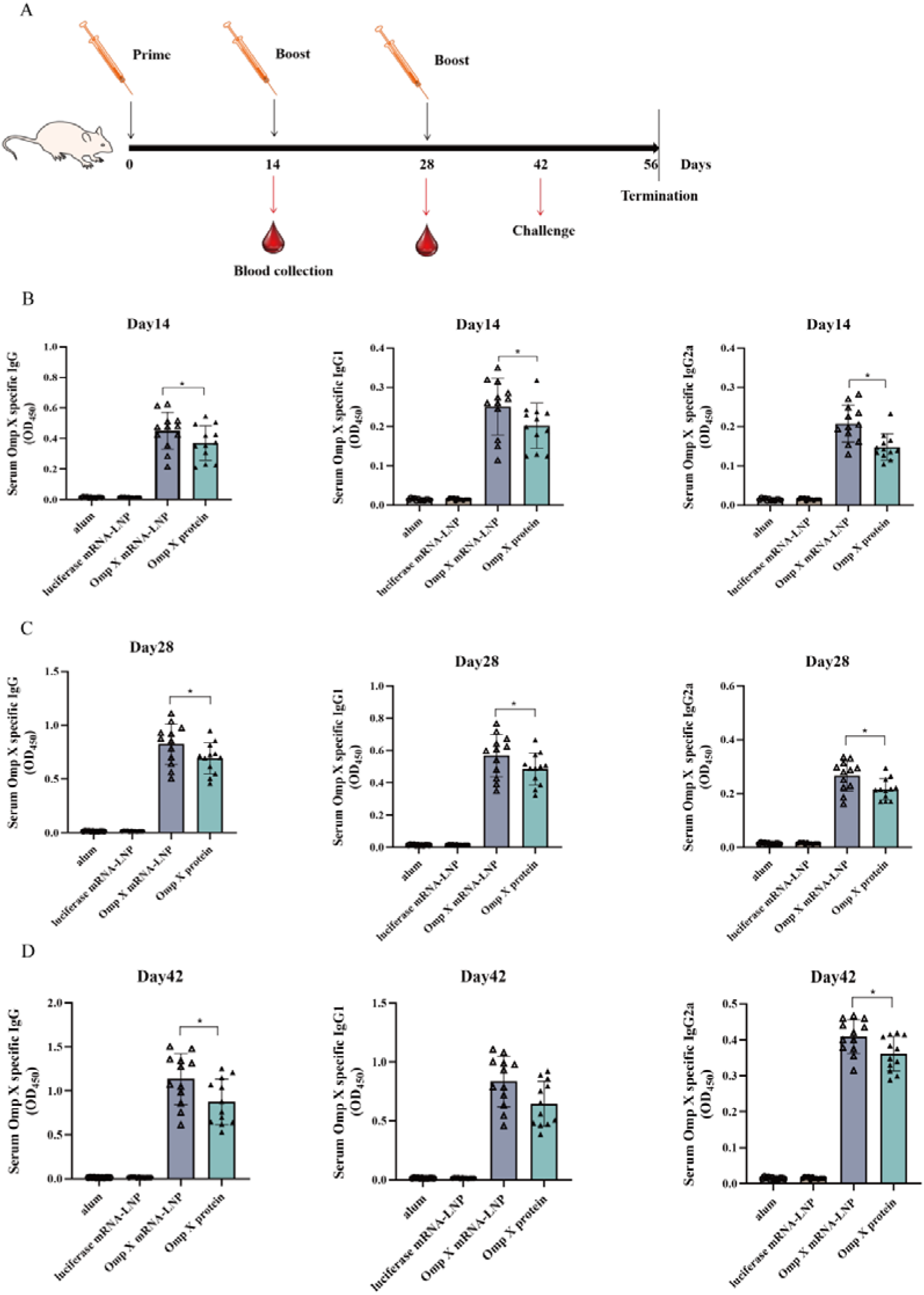
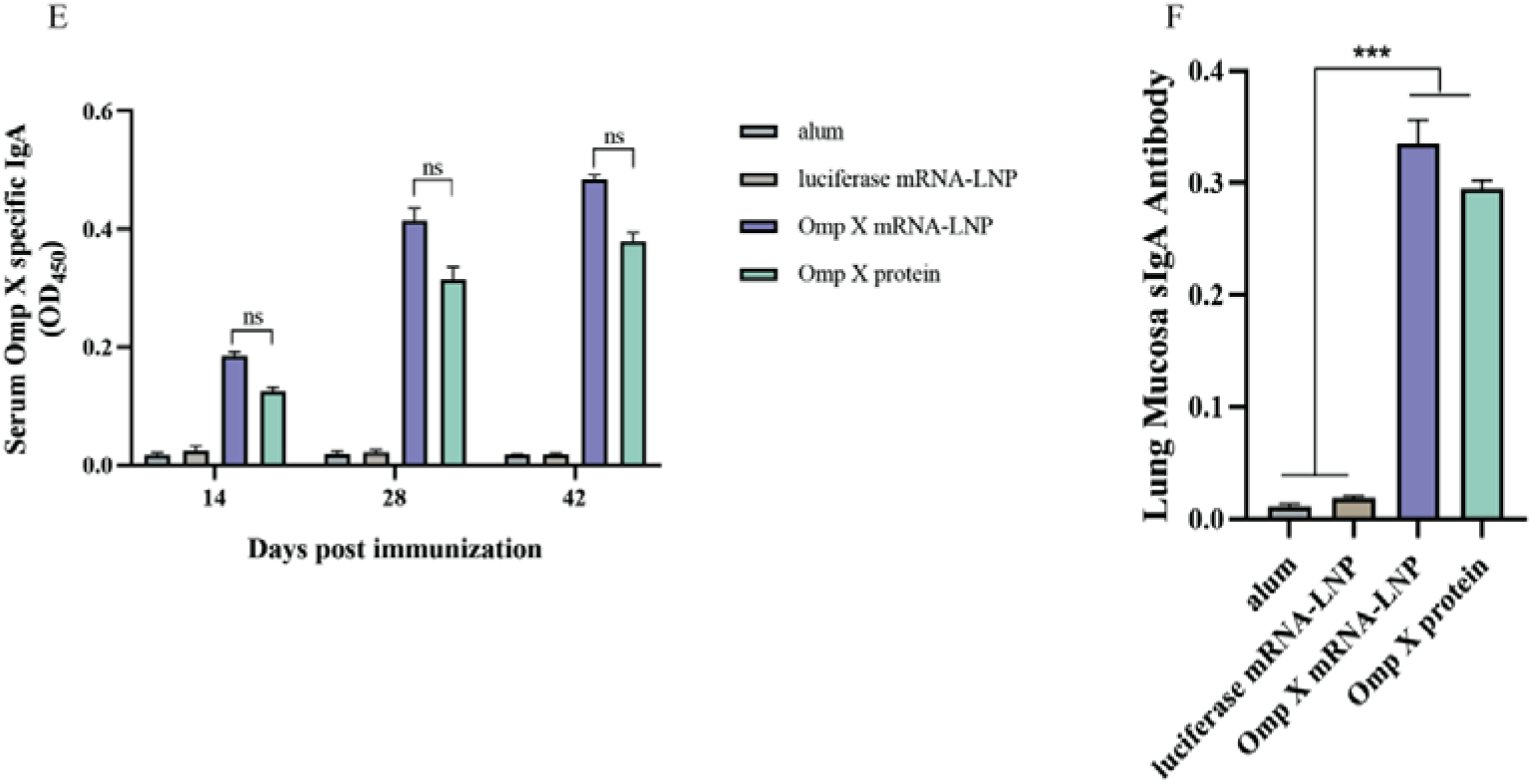
Serum-specific IgG, IgG1, and IgG2a ELISA results in mice. (A) Immunization procedure for the Klebsiella pneumoniae mouse vaccination study; (B) serum OmpX-specific IgG, IgG1, and IgG2a ELISA results on day 14; (C) results on day 28; (D) results on day 42; (E) serum OmpX-specific IgA ELISA results on days 14, 28, and 42; (F) OmpX-specific sIgA ELISA results in lung mucosa.

Immunization significantly elevated IgA antibody levels in the OmpX mRNA vaccine and OmpX protein groups on day 28, peaking on day 42, compared with the alum and luciferase mRNA-LNP groups. Furthermore, it significantly increased the titer of lung mucosal sIgA in the OmpX mRNA vaccine and OmpX protein groups (Fig. 8E, F).

Immunization with mRNA-LNPs and protein vaccines promoted the secretion of IFN-γ, IL-17A, and IL-4 by splenic lymphocytes. IFN-γ and IL-4 were significantly elevated, whereas IL-17A secretion was relatively weak (Fig. 9A-C). Overall, these data suggest that mRNA-LNP vaccination elicits a strong cellular immune response.

**Fig. 9.**
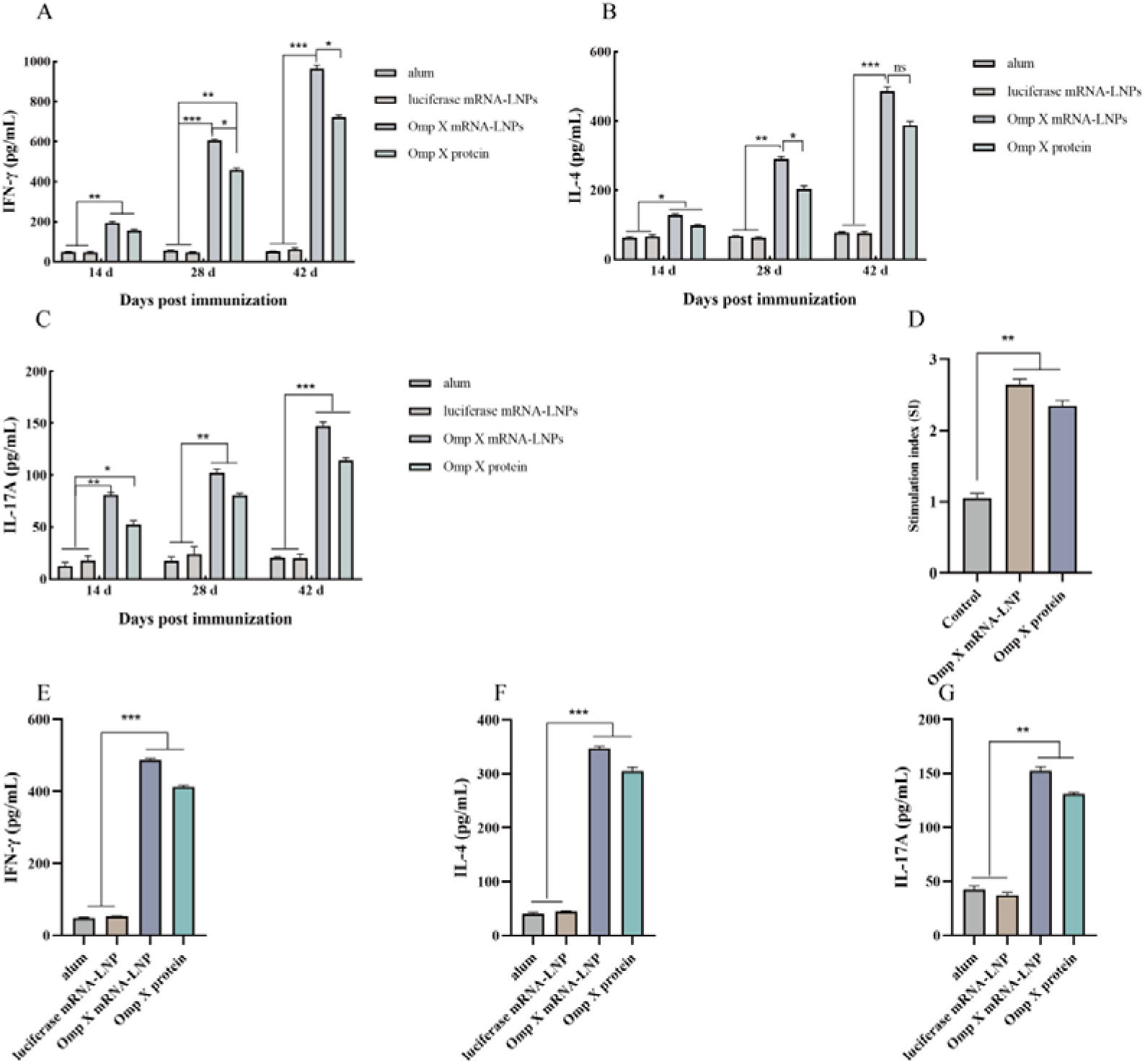
Serum IFN-γ, IL-4, and IL-17A ELISA results and splenic lymphocyte proliferation and cytokine assays in mice. (A) Serum IFN-γ ELISA results; (B) serum IL-4 ELISA results; (C) serum IL-17A ELISA results; (D) OmpX-specific lymphocyte proliferation assay; (E–G) IFN-γ, IL-4, and IL-17A secretion by lymphocytes after restimulation with OmpX protein. The cell culture supernatant was collected for IFN-γ, IL-4, and IL-17A detection by ELISA.

To evaluate antigen-specific T-cell responses, peripheral blood lymphocytes were restimulated with recombinant OmpX protein *in vitro*. The stimulation index was significantly higher in the OmpX-immunized group than in controls (Fig. 9D). IFN-γ, IL-4, and IL-17A secretion by restimulated lymphocytes was measured by ELISA (Fig. 9E-G).

### OmpX-specific T-cell responses and intracellular cytokine expression

mRNA-LNP vaccination significantly increased the frequency of CD4□ and CD8□ T lymphocytes at 42 days post-immunization (Fig. 10A, B). Moreover, mRNA-LNP immunization resulted in greater T-lymphocyte proliferation compared with the protein vaccine group.

**Fig. 10.**
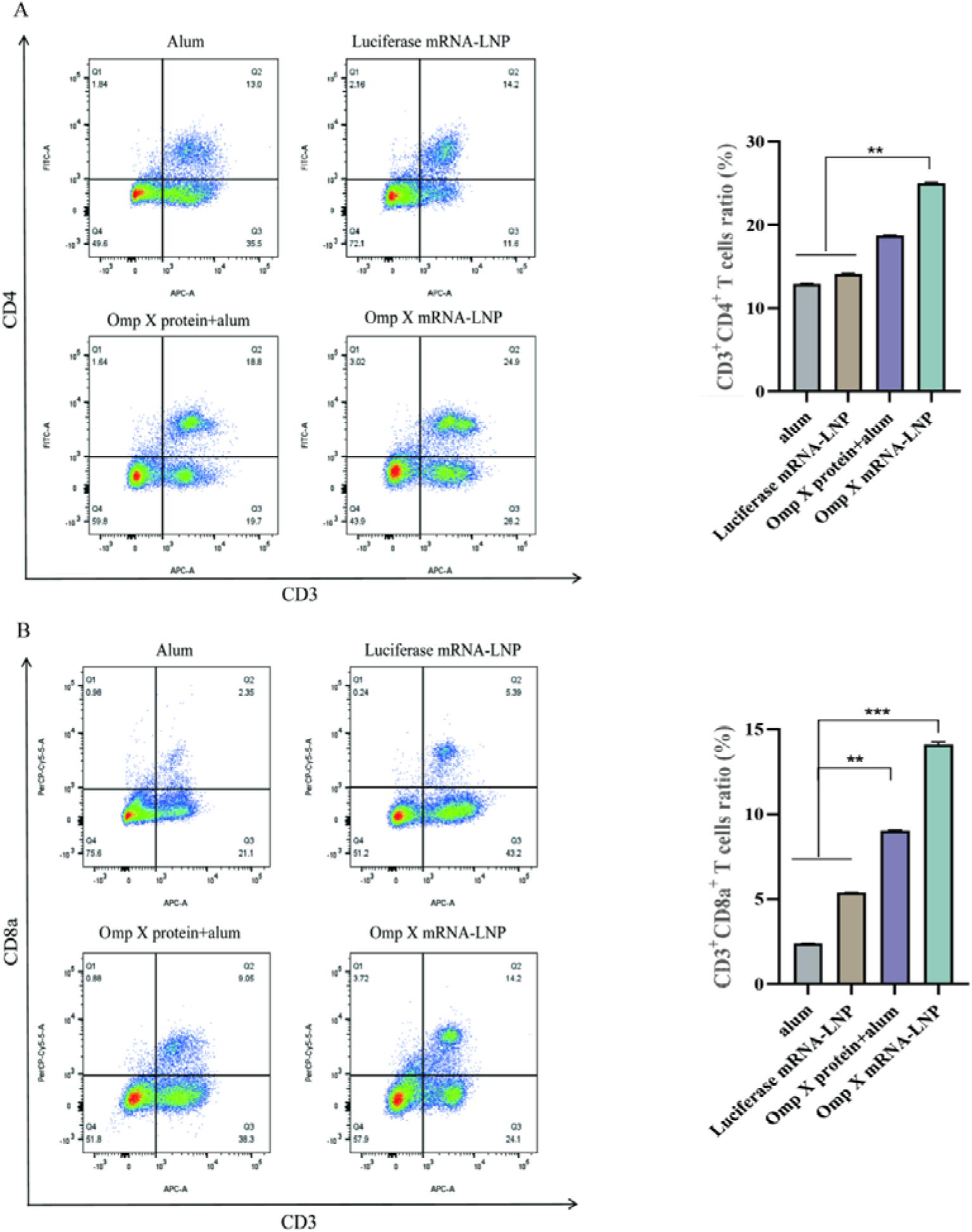
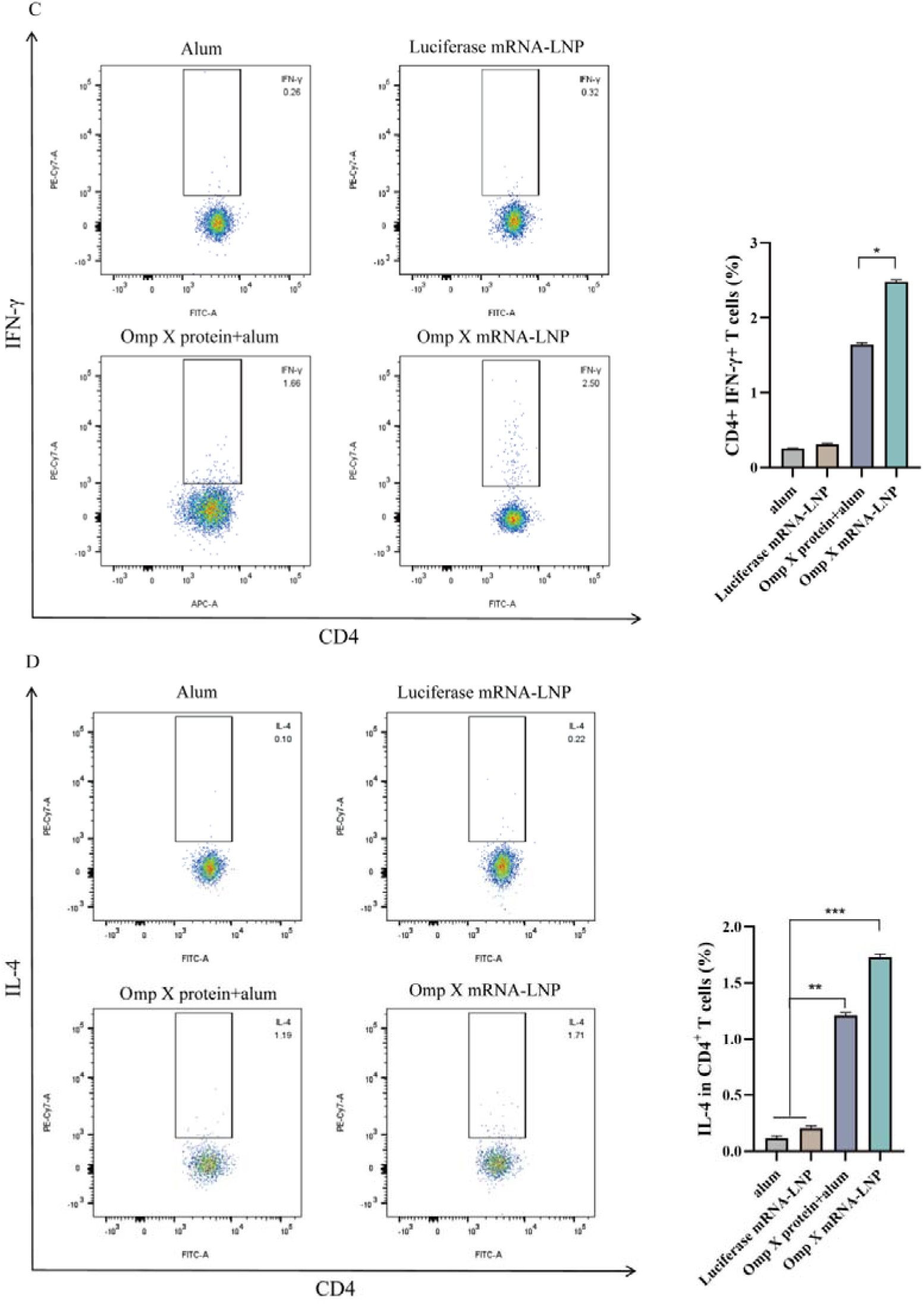
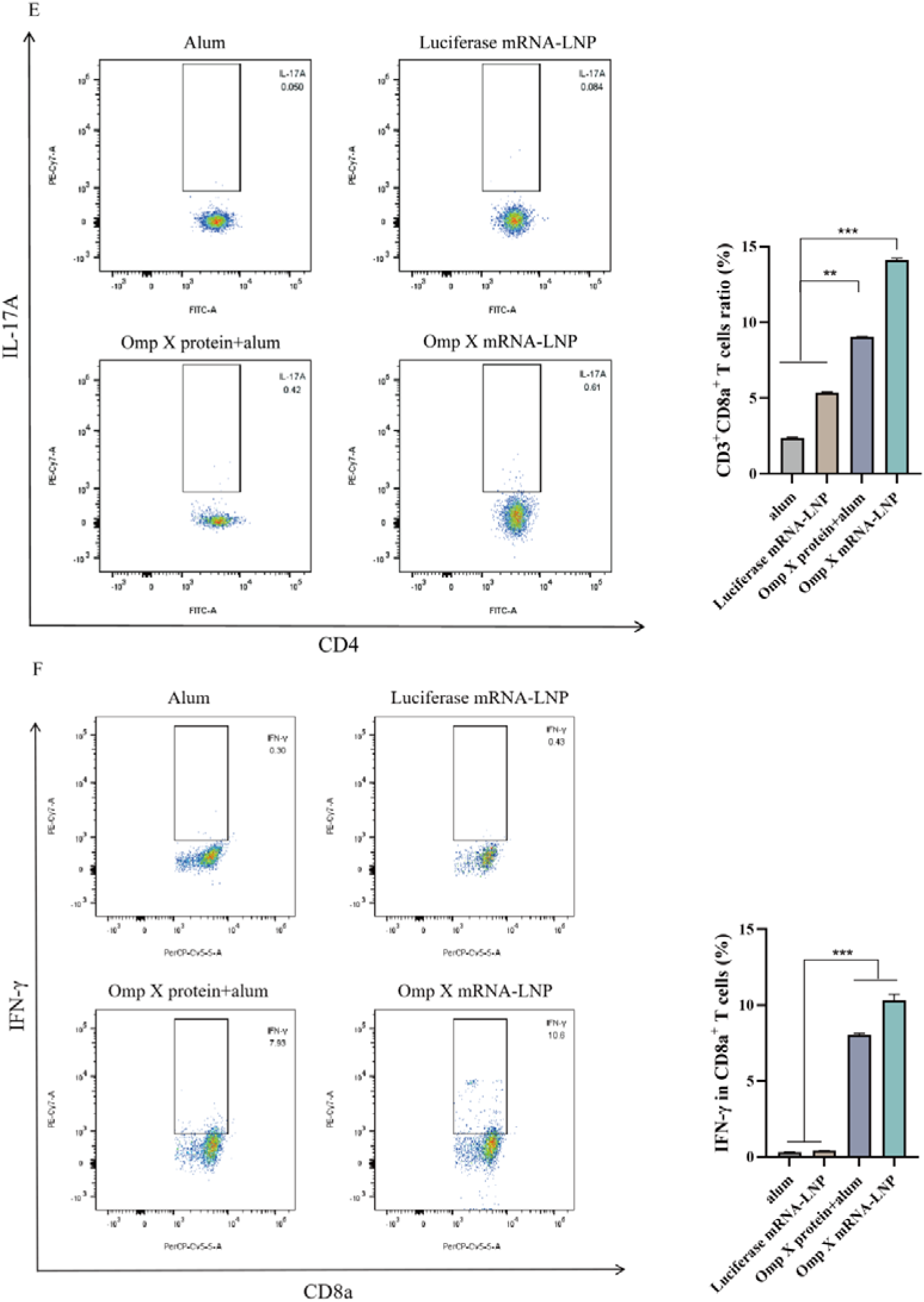
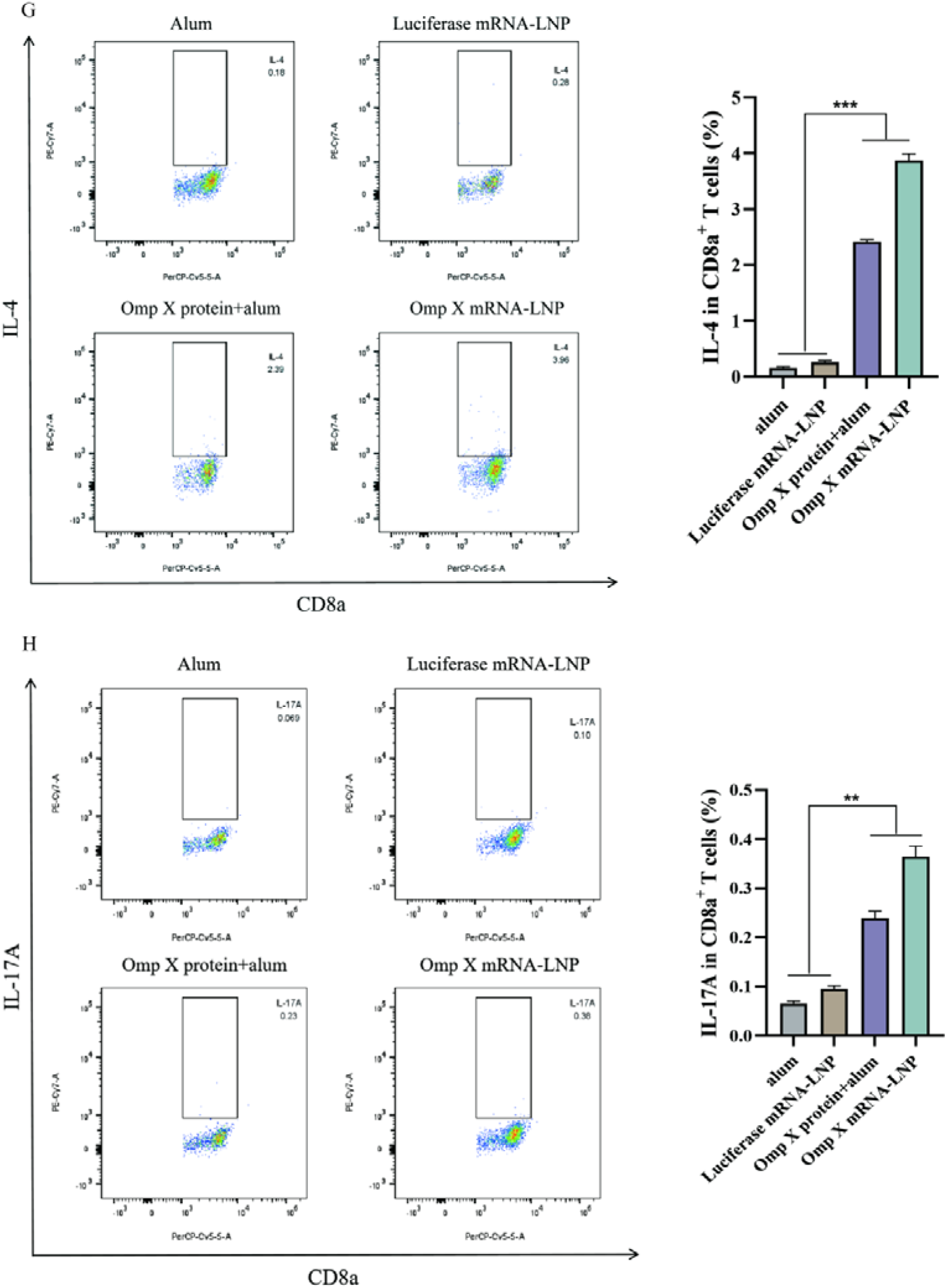
OmpX mRNA-LNP vaccine-induced T-cell responses and expression of the specific cytokines IFN-γ, IL-4, and IL-17A. (A) Flow cytometric analysis of CD3⁺CD4⁺T cells; (B) flow cytometric analysis of CD3⁺CD8a⁺T cells; (C) flow cytometric analysis of IFN-γ⁺CD4⁺T cells; (D) flow cytometric analysis of IL-4⁺CD4⁺T cells; (E) flow cytometric analysis of IL-17A⁺CD4⁺T cells; (F) flow cytometric analysis of IFN-γ⁺CD8a⁺T cells; (G) flow cytometric analysis of IL-4⁺CD8a⁺T cells; (H) flow cytometric analysis of IL-17A⁺CD8a⁺T cells.

The frequencies of IFN-γ□ and IL-4□ T lymphocytes were significantly increased in the mRNA-LNP group compared with controls at 42 days post-immunization (Fig. 10C-H). In contrast, IL-17A□ T-lymphocyte frequencies did not change significantly.

### OmpX-Fc mRNA-LNPs induce protective immunity against KP challenge *in vivo*

The immunization and challenge protocol is shown in Fig. 11A. OmpX-Fc mRNA-LNP immunization significantly reduced bacterial burden in the liver, spleen, lungs, and kidneys compared with the alum and luciferase mRNA-LNP control groups (Fig. 11B, C). Thus, OmpX-Fc mRNA-LNP immunization provides robust protection against systemic KP infection.

**Fig. 11.**
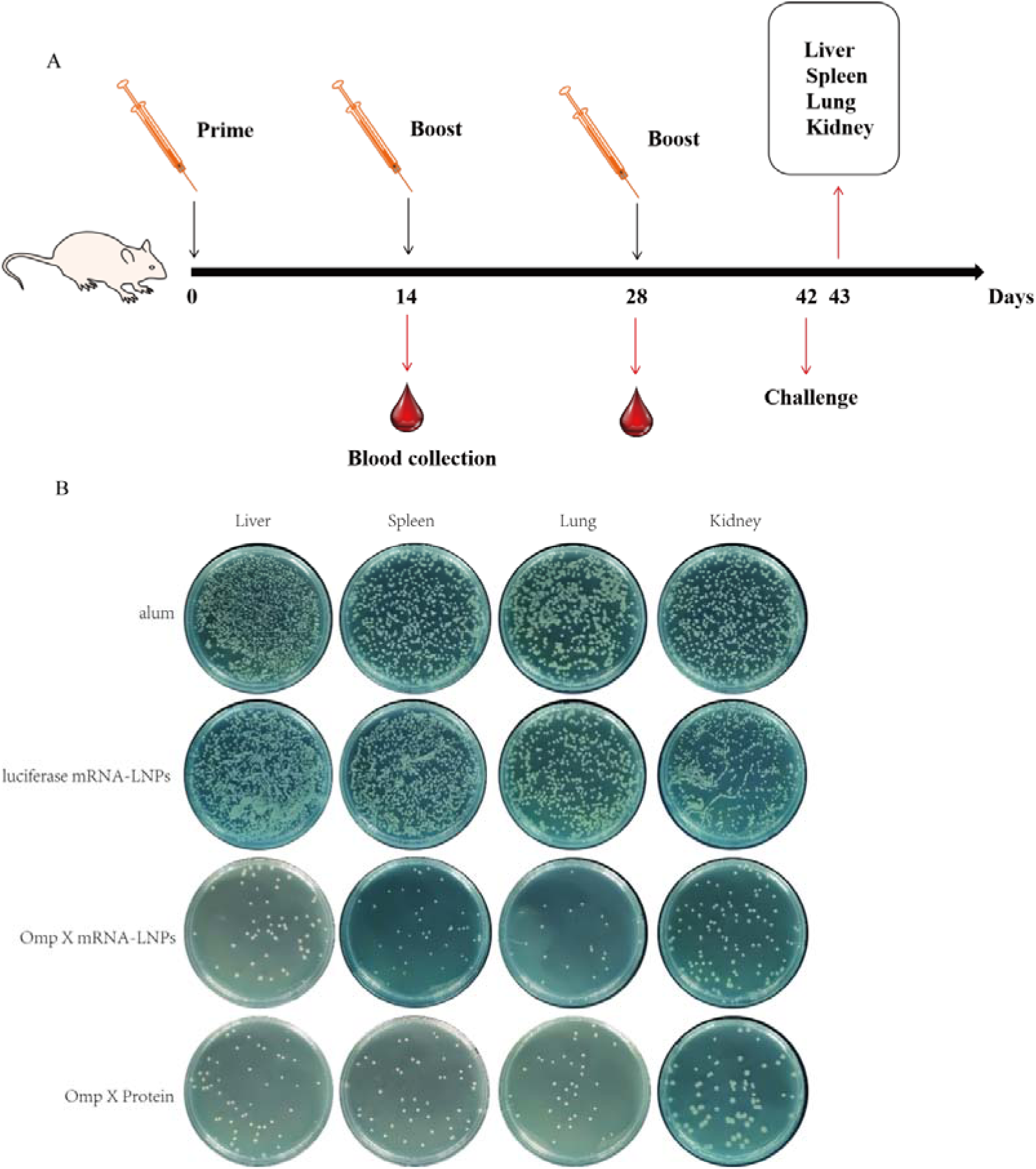
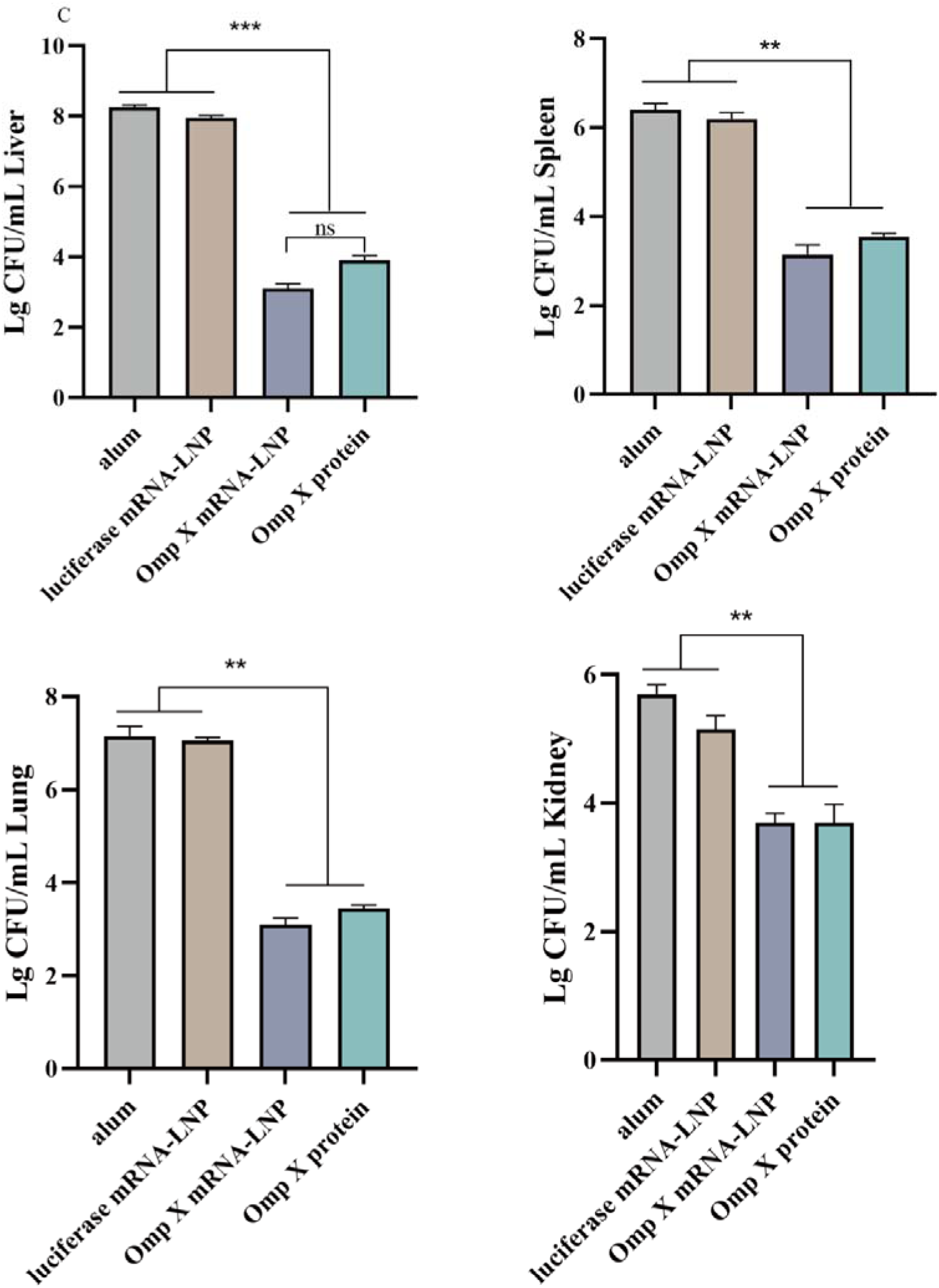
Residual CFU counts in the liver, spleen, lungs, and kidneys after Klebsiella pneumoniae infection in each immunized group of mice. (A) Schematic diagram of the immunization and challenge experiments; (B) single-cell suspensions of liver, spleen, lungs, and kidneys from each immunization group after KP infection incubated on LB plates; (C) statistical results of residual CFU counts in the livers, spleens, lungs, and kidneys of mice in each group.

Vaccinated mice showed significantly greater protection against bacterial dissemination and a marked reduction in splenic CFU compared with controls, indicating a key role for B cells and antibodies in preventing bacterial spread from the lungs. We also observed significant increases in CD4□ and CD8□ T cells in immunized mice after KP challenge, consistent with Th1/Th2-mediated protection.

### The mRNA vaccine protects mice against lethal KP infection

Fig. 12A shows the immunization and challenge schedule. The survival rates of mice immunized with the mRNA vaccine and OmpX protein were 100% and 66.7%, respectively (Fig. 12B). All mice in the alum and luciferase mRNA-LNP groups died within 3 days after challenge. Mice in the mRNA vaccine and OmpX protein groups showed no significant changes in body weight (Fig. 12C), whereas mice in the control groups exhibited significant weight loss after challenge.

**Fig. 12.**
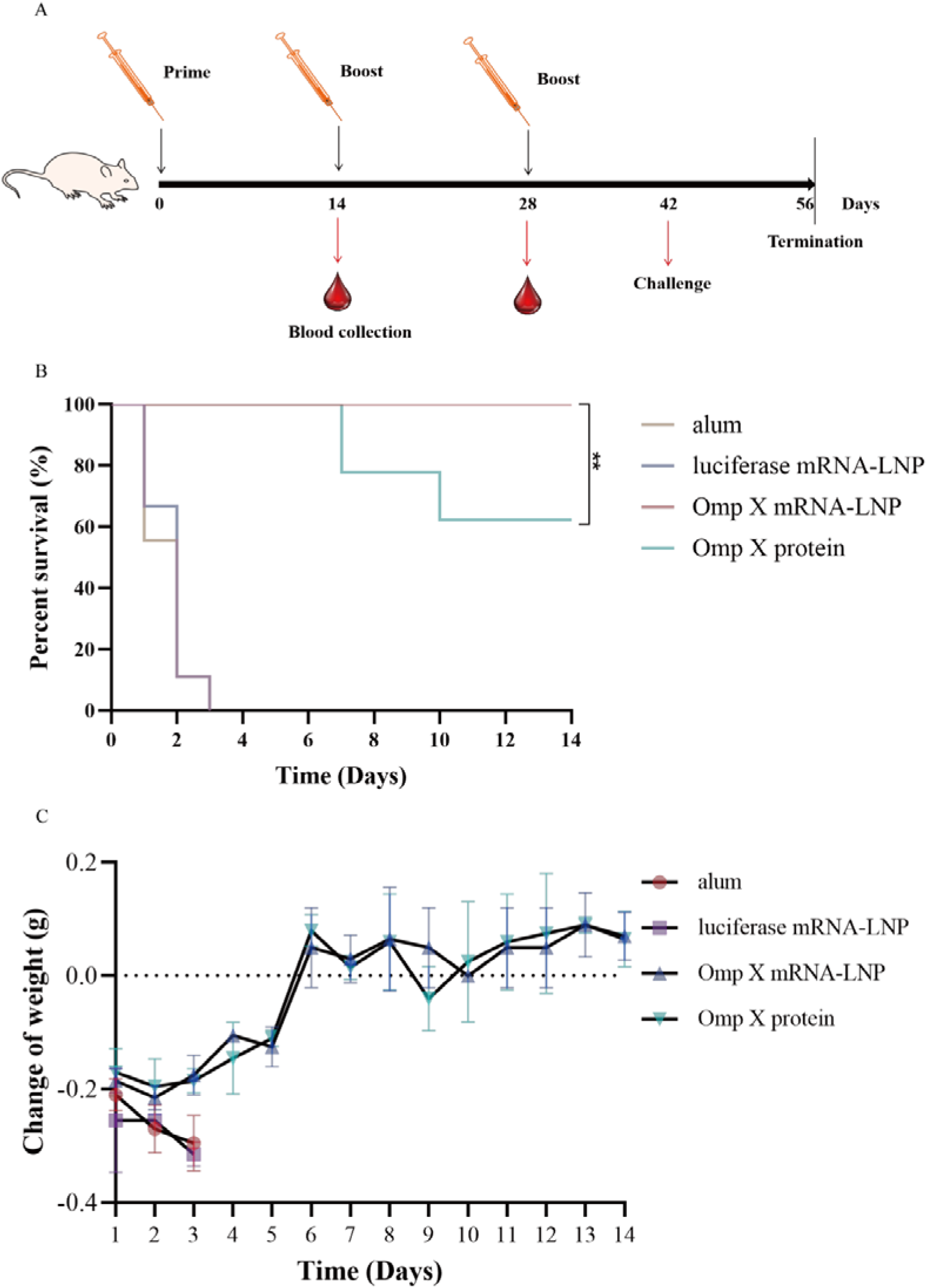
Survival rates and body weight changes of each group of immunized mice after challenge. (A) Schematic diagram of the immunization and challenge experiments; (B) survival curves after challenge in each group of immunized mice; (C) body weight change curves of mice in each group within 14 days after challenge. n=6 mice per group; P values by log-rank (Mantel-Cox) test: mRNA vs alum *P*<0.0001; protein vs alum *P*<0.01.

### Safety profile of the mRNA vaccine

The heart, lung, liver, spleen, and kidney of mice in the alum and luciferase mRNA-LNP groups exhibited necrotic lesions compared with those in the OmpX-Fc mRNA-LNP and OmpX protein groups (Fig. 13A). Mice vaccinated with OmpX-Fc mRNA-LNPs or OmpX protein showed a significant reduction in bacterial load during KP infection, leading to clearance of the invading bacteria and attenuated tissue damage in the lungs and other organs. There were no significant differences in liver and kidney function between treated and control mice (Fig. 13B).

**Fig. 13.**
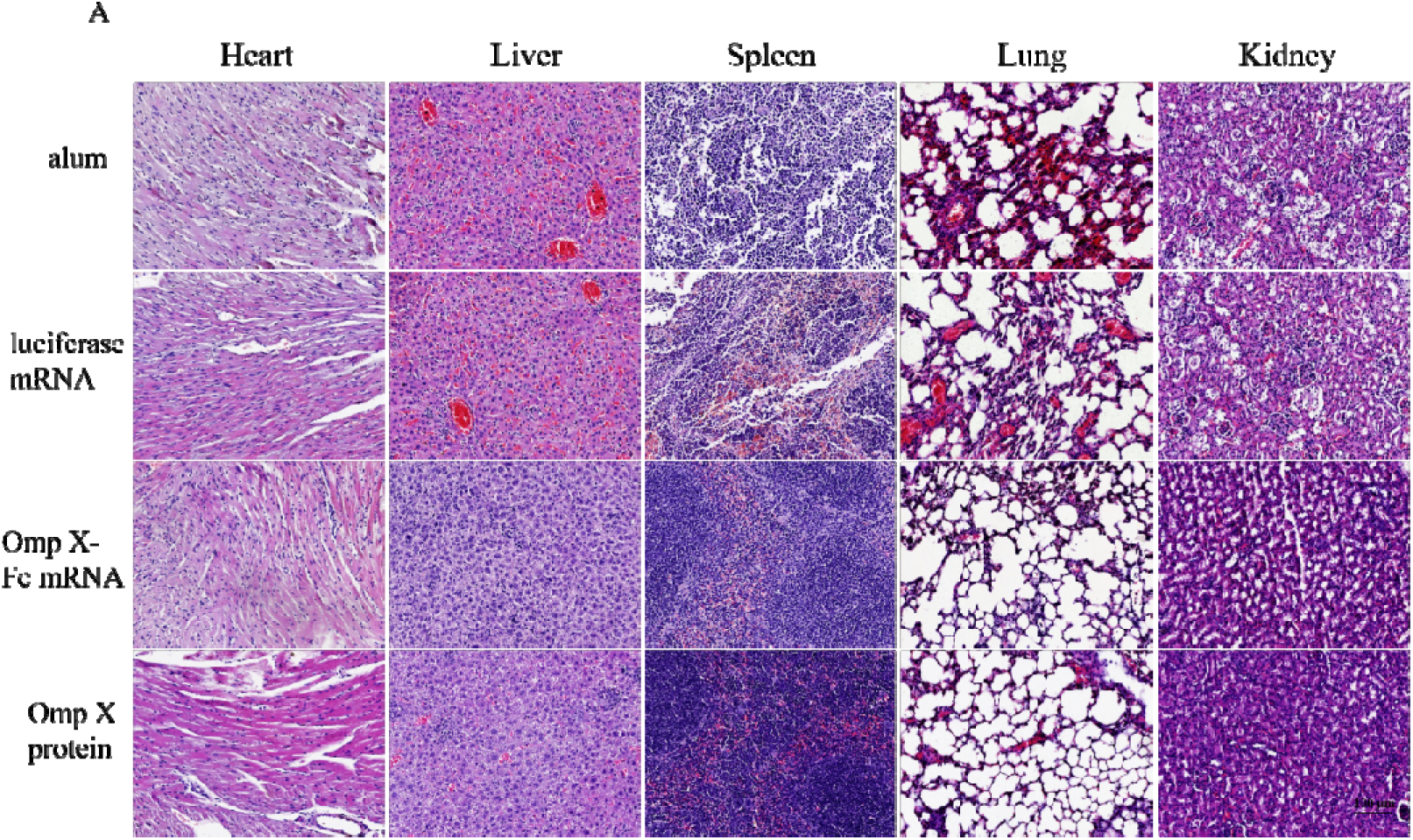
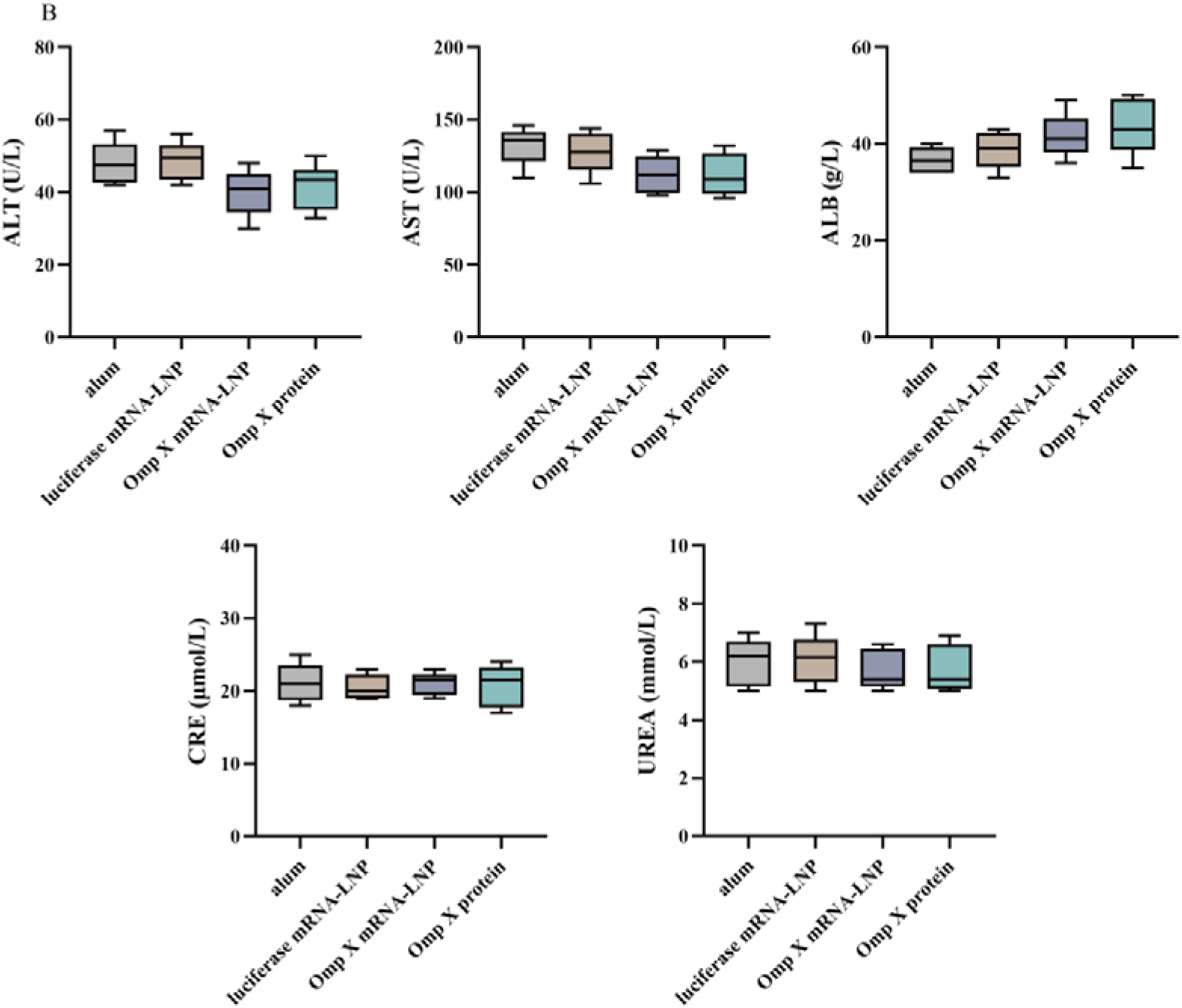
Safety assessment of mRNA-vaccinated mice. (A) H&E staining (scale bar, 100 μm); (B) liver and kidney function measurements in each group of mice.

### OmpX-specific antibodies induce serum bactericidal activity against KP strains

Complement activation is a key component of antimicrobial innate immunity. Mouse anti-OmpX serum from the OmpX mRNA-LNP group achieved average KP lysis rates of 65%, 93%, and 96% at 30, 60, and 90 min, respectively. In comparison, serum from the OmpX protein-immunized group showed lysis rates of 60%, 86%, and 90% at the same time points (Fig. 14). Sera from the alum and luciferase mRNA-LNP control groups failed to induce KP lysis even after 90 min. These results indicate that OmpX-specific antibodies exert direct serum bactericidal activity, which may complement other effector mechanisms in vivo. No lysis was observed with complement alone or with EDTA-treated serum, confirming that killing was complement-dependent.

**Fig. 14.**
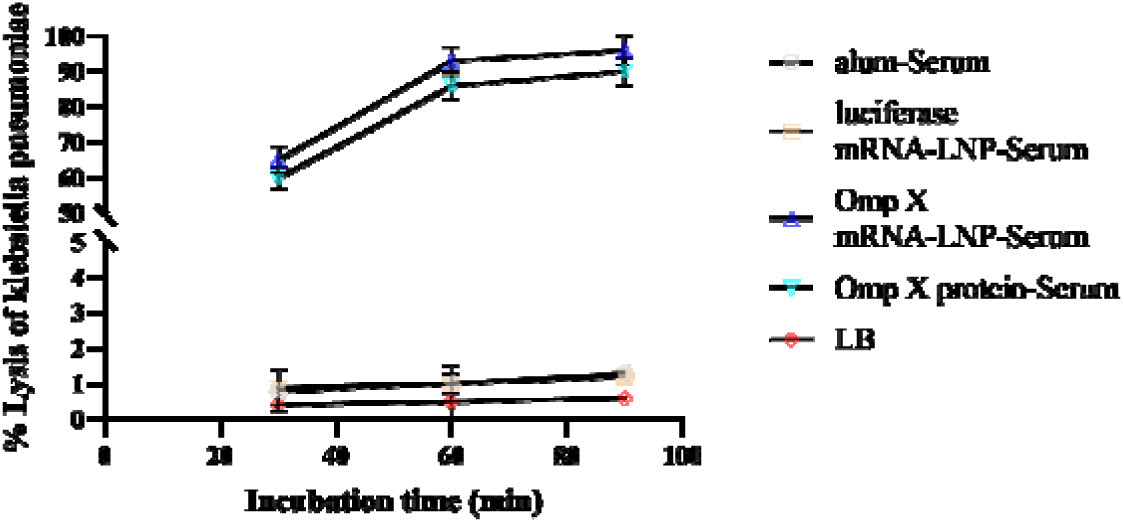
Complement-mediated lysis of KP strains induced by OmpX-specific antibodies.

## DISCUSSION

KP is a leading cause of hospital-acquired pneumonia, with high morbidity and mortality in immunocompromised individuals, including transplant recipients and patients with hematologic malignancies infected with MDR strains. As a clinically important pathogen, KP is associated with severe pneumonia, meningitis, sepsis, and other nosocomial infections [16, 17]. KP infections impose a substantial global societal burden and healthcare cost. The emergence and spread of antibiotic resistance have increasingly limited therapeutic options, particularly in hospital settings. Moreover, the complex pathogenicity and strain heterogeneity of KP have hindered the development of universally protective vaccines. Novel and effective vaccines are therefore essential for controlling KP infections.

This study evaluated the potential of an OmpX-based vaccine candidate through integrated bioinformatics and experimental validation. Bioinformatics analyses confirmed that OmpX is a highly conserved protein widely distributed across different KP strains. The predicted secondary structure contains abundant β-strands and random coils, with numerous exposed regions suitable for antigenic epitope formation. The antigenicity score of OmpX was 1.01. Both OmpX mRNA and OmpX protein immunization significantly elevated OmpX-specific IgG, IgG1, and IgG2a levels in mice. Humoral immunity plays a critical role in eliminating bacterial infections. The high IgG titers in OmpX antiserum induced complement-mediated KP lysis via the classical pathway. Porin proteins from KP can form complexes with C1q [18, 19, 43], a component of the classical complement pathway, thereby activating the complement cascade—a property not shared by other outer membrane proteins. Activation of the alternative complement pathway has also been reported as an efficient mechanism for eliminating serum-sensitive KP strains, which may partly explain the high protection observed in immunized mice. Additionally, OmpX-induced serum antibodies may functionally prevent bacterial adhesion, while opsonizing antibodies promote macrophage-mediated killing and complement activation. Future studies should investigate the mechanisms of OmpX immunogenicity in macrophage adhesion and the protective role of serum antibodies, which may be particularly important for individuals with impaired cellular immunity or severe inflammatory conditions in the late stages of bacteremia. OmpX immunization induced a robust systemic antibody response, and the antibodies were intrinsically protective in vitro, contributing meaningfully to protection against KP infection.

In addition, splenocytes from mice immunized with the OmpX mRNA or OmpX protein vaccines proliferated vigorously in vitro and secreted substantial amounts of IFN-γ, IL-4. OmpX triggered Th1- and Th2-type cellular immunity, activating T-helper cell subsets. Cytotoxic T lymphocytes also contributed to KP clearance, aiding protection against infection. OmpX immunization protected mice from subsequent bacterial challenge and significantly reduced KP-induced pneumonia and hemorrhage in visceral organs. The protective efficacy of the OmpX-based vaccine can be attributed to several factors. First, OmpX immunization induced higher IL-4 levels than controls; both Th1 (IFN-γ, IgG2a) and Th2 (IL-4, IgG1) responses were induced; in mice, complement-fixing IgG2a antibodies likely drive the observed serum bactericidal activity via the classical pathway, consistent with the ability of KP porins to bind C1q [19]. Second, KP significantly induces memory T cells [20, 22]. In this study, three doses of OmpX mRNA-LNPs induced effective memory reactivation upon challenge. Finally, the route of vaccination strongly influences the immunogenicity of different vaccine types [23, 24]. Overall, vaccine efficacy is closely related to antigen type, dosage, and administration route [25–27].

In conclusion, this study clearly demonstrates that the OmpX mRNA vaccine induces a robust adaptive immune response and protects immunized mice against KP infection. Notably, OmpX mRNA immunization significantly improved survival following KP challenge, achieving a 100% survival rate. These findings establish OmpX mRNA-LNPs as an effective platform for developing next-generation KP vaccines. Future studies should evaluate this vaccine strategy in other mammalian models to further advance the fight against KP-associated diseases.

## Conclusion

Integrated bioinformatics and empirical validation identified KP OmpX-a conserved, surface-exposed outer membrane protein-as a potent vaccine antigen conferring immunoprotection in murine models. Our findings demonstrate that the OmpX mRNA-LNP vaccine elicits robust Th1/Th2-polarized humoral and cellular immunity and pathogen-neutralizing antibodies, achieving 100% survival and significant reductions in bacterial burden across target organs post-challenge. These data position OmpX as a promising candidate for next-generation KP vaccines.

## CRediT authorship contribution statement

**Hongxian Chen**: Writing-review & editing, **Yunwang Zhao**: Writing-original draft, **Shiyi Zhou**: Visualization, **Lei He and Chenlin Xiu**: Methodology, Formal analysis, **Baihai Huang**: Conceptualization. **Xiongxiong Lu**: Writing-review & editing, Writing-original draft, Methodology, Conceptualization.

## Acknowledgments

This work was supported in part by the National Natural Science Foundation of China (81560346).

## Availability of data and materials

The datasets used during the current study are available from the corresponding author upon reasonable request.

## Declarations

### Ethics approval and consent to participate

All mouse immunizations were performed in accordance with the ethical regulations for animal testing and research.

---

**Supplementary Fig. S1.**
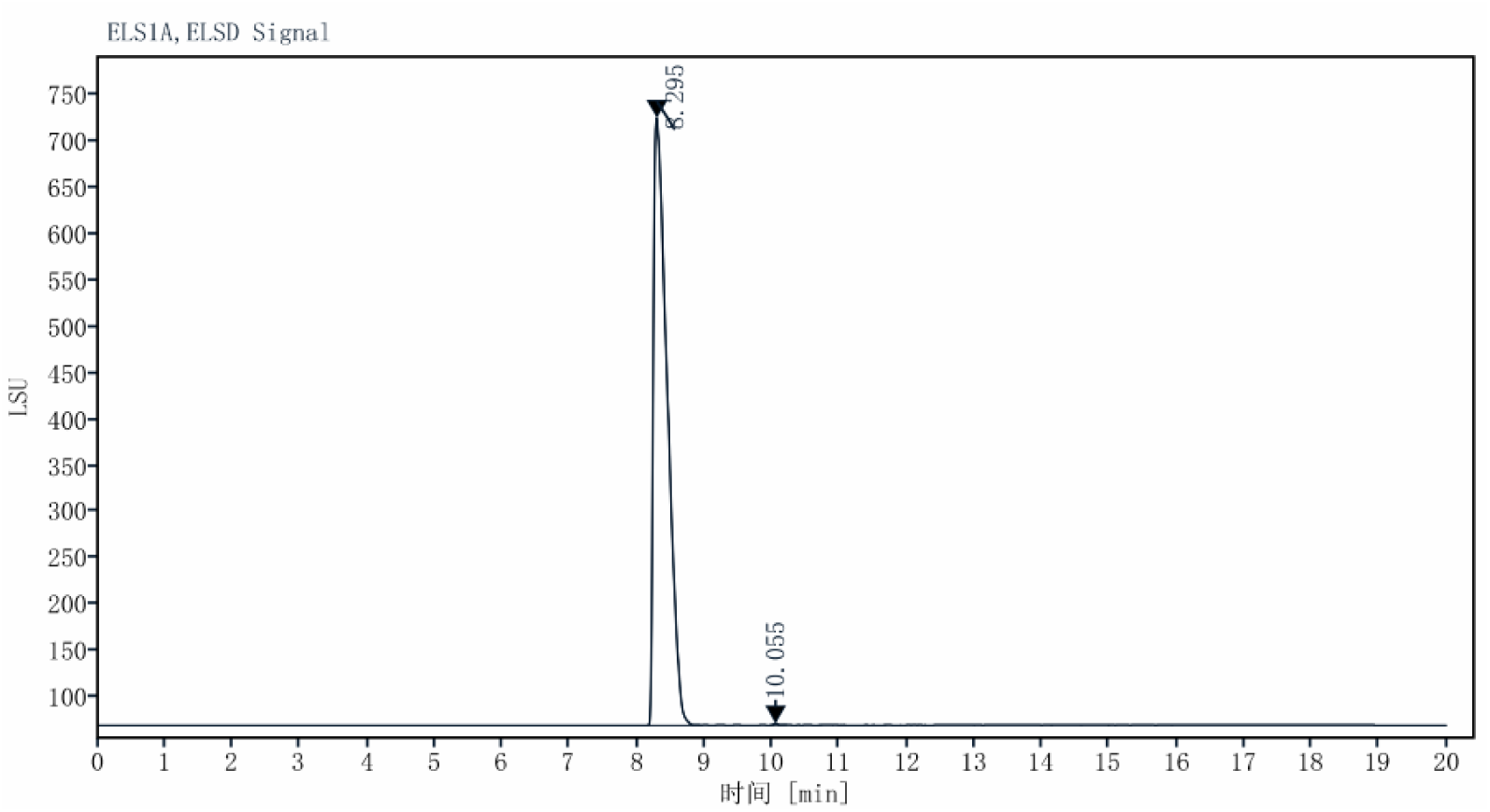
Mass spectra of the ionizable lipid molecules.

**Supplementary Figure S2.**
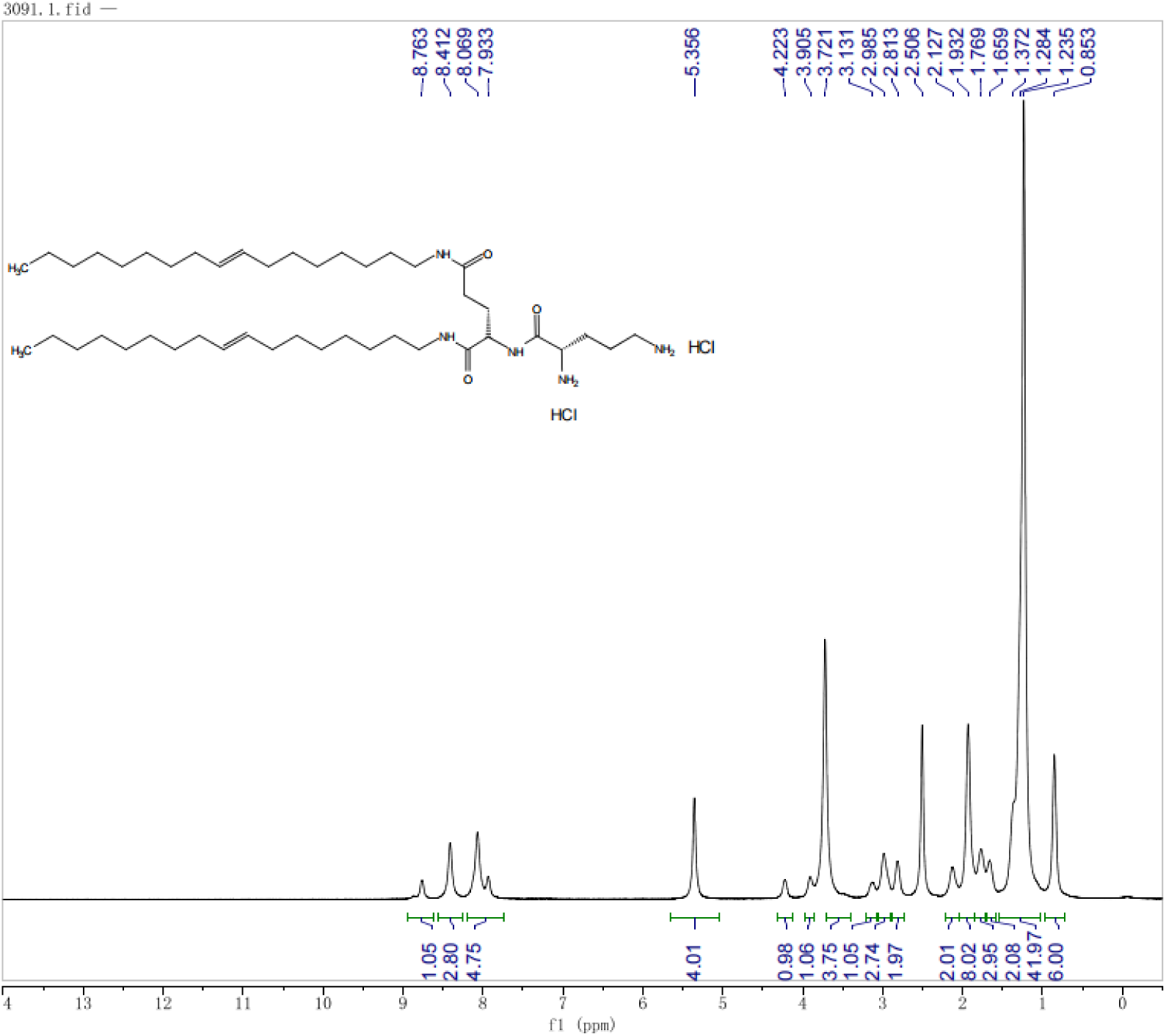
¹H NMR spectrum of the ionizable lipid molecules.

